# Bioorthogonally Cross-Linked Hyaluronan-Laminin Hydrogels for 3D Neuronal Cell Culture and Biofabrication

**DOI:** 10.1101/2021.09.27.461549

**Authors:** Michael Jury, Isabelle Matthiesen, Fatemeh Rasti Boroojeni, Saskia L. Ludwig, Livia Civitelli, Thomas E. Winkler, Robert Selegård, Anna Herland, Daniel Aili

**Affiliations:** Laboratory of Molecular Materials, Division of Biophysics and Bioengineering, Department of Physics, Chemistry and Biology, Linköping University, 581 83 Linköping, Sweden; Division of Micro and Nanosystems, KTH Royal Institute of Technology, 10044 Stockholm, Sweden; Institute of Microtechnology & Center of Pharmaceutical Engineering, Technische Universität Braunschweig, 38106 Braunschweig, Germany; AIMES, Center for Integrated Medical and Engineering Science, Department of Neuroscience, Karolinska Institute, 17165 Solna, Sweden; Nuffield Department of Clinical Neurosciences, John Radcliffe Hospital, West Wing, University of Oxford, Oxford OX3 9DU, United Kingdom

**Author notes:** M. Jury and I. Matthiesen contributed equally to this work.

**Keywords:** Hyaluronan, hydrogel, laminin, neural stem cells, 3D cell culture, 3D bioprinting

## Abstract

Laminins (LNs) are key components in the extracellular matrix of neuronal tissues in the developing brain and neural stem cell niches. LN-presenting hydrogels can provide a biologically relevant matrix for the 3D culture of neurons towards development of advanced tissue models and cell-based therapies for the treatment of neurological disorders. Biologically derived hydrogels are rich in fragmented LN and are poorly defined concerning composition, which hampers clinical translation. Engineered hydrogels require elaborate and often cytotoxic chemistries for cross-linking and LN conjugation and provide limited possibilities to tailor the properties of the materials. Here we show a modular hydrogel system for neural 3D cell culture, based on hyaluronan (HA) and poly(ethylene glycol) (PEG), that is cross-linked and functionalized with human recombinant LN 521 using bioorthogonal copper-free click chemistry. Encapsulated human neuroblastoma cells demonstrate high viability and grow into spheroids. Neuroepithelial stem cells (lt-NES) cultured in the hydrogels can undergo spontaneous differentiation to neural fate and demonstrate significantly higher viability than cells cultured without LN. The hydrogels further support the structural integrity of 3D bioprinted structures and maintain high viability of syringe extruded lt-NES, which can facilitate the development of advanced neuronal tissue and disease models and translation of stem cell-based therapies.

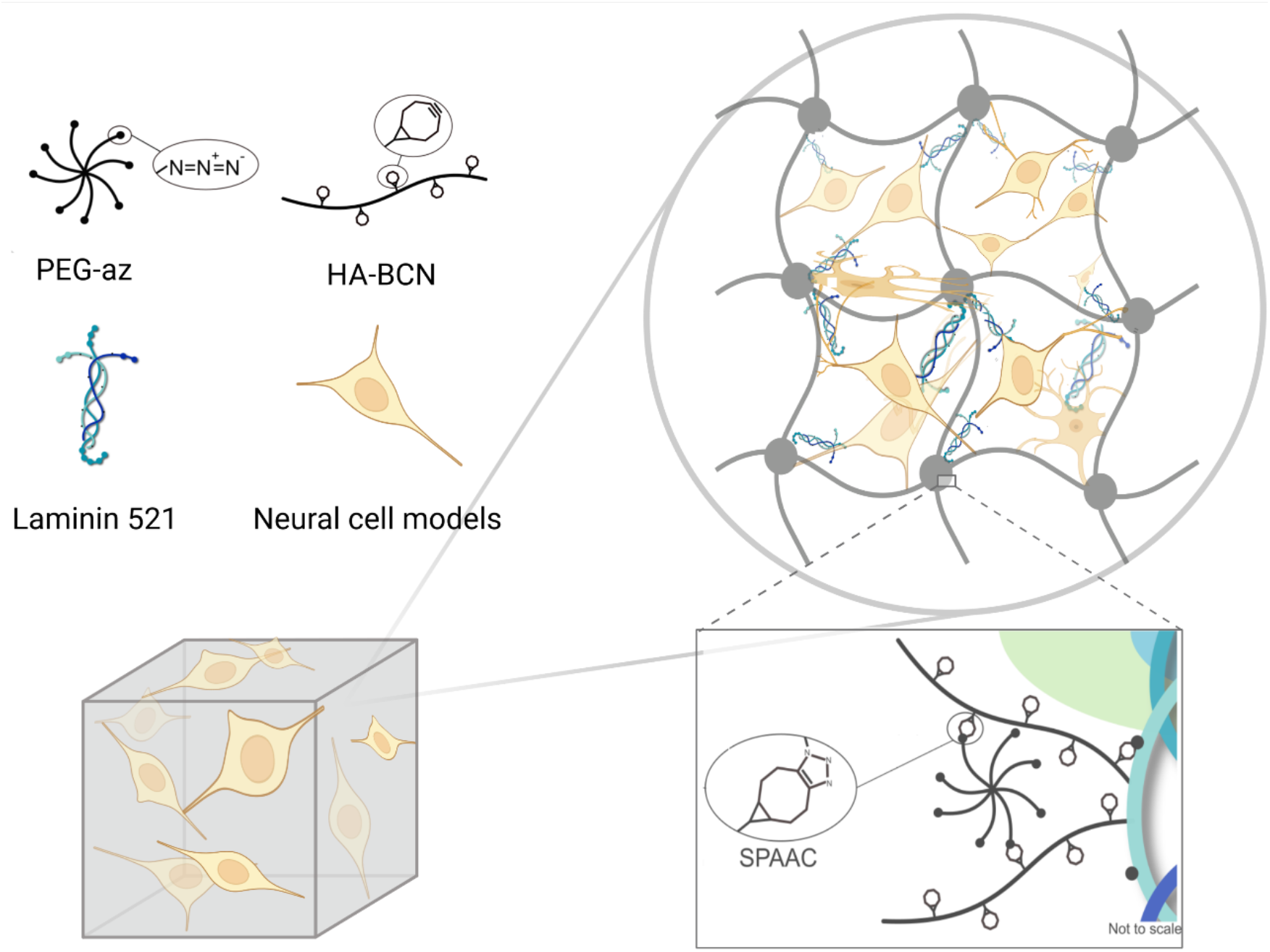
The authors present an extracellular matrix mimicking hydrogel for 3D culture of neural cell models. Based on hyaluronic acid and poly(ethylene glycol), the hydrogel immobilizes recombinant laminin 521, associated with neuronal development. The study demonstrates support of neuroblastoma cell viability, spontaneous human neuroepithelial stem cell differentiation, and the protective effect of the hydrogels during bioprinting and syringe needle ejection.

## Introduction

Neurological disorders caused by tumors, degeneration, trauma, infections, congenital or structural defects are combined the second leading cause of death globally.^[1,2]^ Access to physiologically relevant human neuronal tissue- and disease models is required to improve treatment outcomes and accelerate drug development, which has sparked considerable interest in techniques for generating organoids, organs-on-chips and 3D bioprinted constructs with tissue- and organ-like properties.^[3–5]^ New innovative technologies have further facilitated this development for additive manufacturing and advancements in stem cell technologies.^[6]^ The latter has also spawned many opportunities for exploring and translating novel therapeutic strategies for neurodegenerative disorders or traumatic injuries.^[7–10]^ Due to the high sensitivity of neural tissues to damage and their minimal regenerative capacity, reparative and regenerative treatment modalities based on stem cell transplantation offer new possibilities to relieve symptoms and restore function after injury or disease.^[11,12]^ Both the engineering of functional cellular architectures and the development of cell-based therapies require well-defined materials that can mimic the function of the native extracellular matrix (ECM).^[13–15]^ The ECM offers structural support for cells in all tissues and organs, orchestrates numerous cellular processes, and is critical for cell function and guiding cell behavior and differentiational fate.^[16,17]^ The ECM is a dynamic and spatially heterogeneous biomolecular material comprised of glycosaminoglycans (GAGs), proteoglycans, glycoproteins, and fibrous proteins.^[18]^ In neural tissues the ECM has a unique composition with large quantities of lecticans and GAGs, such as hyaluronic acid (HA), whilst collagen, vitronectin, fibronectin and other fibrous proteins are less abundant. ^[19,20]^ HA is critical for neuronal development and commonly localized in neural stem cell niches.^[21,22]^ The developing brain is also rich in laminins (LNs).^[23]^ LNs are large heterotrimeric proteins (400 – 900 kDa), consisting of an α, β, and λ-chain, and are closely associated with neuronal development and known to promote and guide neurite outgrowth^[24]^ and to stabilize neuronal synapses.^[25]^

In addition to providing an adequate and biologically relevant microenvironment, ECM mimicking materials developed for biofabrication, and therapeutic applications must be compatible with the required processing conditions, including syringe extrusion, while maintaining high cell viabilities. Whereas biologically derived hydrogels, such as Matrigel, to a certain extent fulfill the requirements for biological relevance, these animal-derived materials are poorly defined concerning composition and can suffer from large batch-to-batch variations that can compromise reproducibility and make clinical translation very challenging. In addition, the limited possibilities to tailor the properties of biologically derived ECM hydrogels make them difficult to adapt to a 3D bioprinting process or to integrate into microfluidic devices for the development of organ-on-chips. Engineered ECM-mimicking materials are typically designed with the ambition to address these shortcomings. Biopolymers, such as alginate,^[26]^ collagen,^[27]^ elastin,^[28]^ hyaluronic acid,^[29]^ and synthetic polymers based on, e.g., poly(ethylene glycol) (PEG),^[30]^ poly (vinyl alcohol) (PVA),^[31]^ or poly(2-hydroxyethyl methacrylate) (pHEMA),^[32]^ are widely used in the fabrication of ECM mimicking hydrogels. The potential to process the hydrogels and their performance for cell culture is, in addition to composition, highly dependent on the polymer cross-linking chemistry and network topology as well as cross-linking kinetics and density.^[33,34]^ Whereas supramolecular cross-linking strategies based on molecular self-assembly or ion-coordination are well tolerated by cells and allow for in-situ/in-vivo gelation, the resulting hydrogels are inherently weak and dynamic, leading to uncontrolled and gradual dissolution over time.^[35]^ Covalently cross-linked hydrogels are typically more robust and can cover a wider stiffness range but often rely on chemistries that can harm cell viability, such as UV-triggered photo-polymerization or involve reactions that are difficult to control in a biological context due to cross-reactivity or poor stability of the functional groups.^[36]^ Bioorthogonal strategies, e.g., copper-free click chemistry, have emerged as attractive options for hydrogel cross-linking and can facilitate *in situ* cell encapsulation and biofabrication.^[29,37–39]^

Because of the critical role of LNs in neural tissue development,^[40]^ several different strategies have been developed to incorporate full-size LN in engineered ECM mimicking hydrogels to mimic the native 3D microenvironment better. Whereas affinity-based interactions^[41]^ or physical trapping of LN in the hydrogel network^[42]^ reduce the risk of interfering with LN structure and function, the LN gradually dissociates from the hydrogels over time. Covalent conjugation can result in more efficient retention of LN in the hydrogels^[43,44]^ but can compromise LN function if not carefully optimized. Difficulties in controlling and tuning both cross-linking kinetics and LN biofunctionalization simultaneously^[45–47]^ can further complicate the development of generic LN-presenting hydrogel systems for cell-based therapeutics and bioinks.

In this work, we have developed an injectable and 3D bioprinting-compatible modular HA-based hydrogel system that allows for convenient integration and efficient retention of recombinant LN-521. Recombinant LN-521 has been widely used for stem cell expansion and the generation of neural progenitor cells for disease models and stem cell therapies.^[15,48,49]^ We cross-linked the LN-521 presenting hydrogels by bioorthogonal copper-free click chemistry, which enabled tuning of material properties and creation of biologically relevant microenvironments evidenced by supported encapsulation and culture of both human neuroblastoma cells (SH-SY5Y) and human induced pluripotent stem cell (hiPSC)-derived long-term neuroepithelial stem cells (lt-NES). Furthermore, the hydrogels’ favorable and tunable rheological properties provided a protective effect on the lt-NES during syringe extrusion in an *in vitro* model for cell injection therapy. In addition, 3D bioprinting of the cell-laden hydrogels allowed for the fabrication of structurally well-defined constructs with high cell viabilities, facilitating further development of advanced tissue and disease models.

### Experimental Section

Detailed methods can be found in the Supporting Information.

#### Laminin labeling and formation of hydrogel

Hyaluronan-poly(ethylene glycol) (HA:PEG) hybrid hydrogels were prepared by combining bicyclo[6.1.0]nonyne (BCN) modified HA (∼100 kDa) and an 8-arm PEG with terminal azides ((PEG-Az)_8_) as previously described.^[50,51]^ LN was modified with azide (Az) moieties using linkers of different lengths (LN-Az and LN-p-AZ) and was conjugated to HA-BCN, after which (PEG-Az)_8_ was added to form the final hydrogel at 37°C. The hydrogels were analyzed by rheology and scanning electron microscopy (SEM). In addition, the effect of Az-functionalization on LN retention was measured using fluorescence spectroscopy.

#### SH-SY5Y cell culture and differentiation

SH-SY5Y cells, differentiated and undifferentiated, were encapsulated in 1 % and 2 % w/v hydrogels, with and without LN and cultured for 10 days. Cell viability was assessed using an Alamar blue (AB) assay at 3, 7 and 10 days of culture, and stained for the HA-receptor CD44 and actin and imaged using confocal fluorescence microscopy.

#### Encapsulation and spontaneous 3D-differentiation of lt-NES

lt-NES were encapsulated in HA-based LN functionalized hydrogels, cultured for 7 days, and allowed to differentiate spontaneously. The results were benchmarked against both Matrigel and 2D cultures on tissue culture plates. The effects of LN on differentiation and viability were investigated using AB at 1, 3 and 7 days of subculture. After 7 days of subculture, the mRNA expression of the stem cell markers SOX2 and NES and the neuronal markers DCX, TUBB3 and SYN1 were investigated with qPCR. We further stained the hydrogels for DCX and F-actin for confocal imaging to visualize the morphology of the cells including neurite outgrowth.

#### Syringe ejection and 3D bioprinting

To test whether the hydrogel could serve as a protecting matrix for stem cell therapy applications, lt-NES were encapsulated in HA:PEG hydrogels and ejected through a 27G syringe needle using a syringe pump. In comparison, we ejected cells through the syringe needle in their media. After ejection, cell viability was determined both at an immediate stage and after 24h using a LIVE/DEAD stain. SH-SY5Y cells in HA:PEG hydrogels were bioprinted using a Cellink BioX equipped with a 27G needle. Cell viability was investigated using a LIVE/DEAD assay and imaged using confocal microscopy.

#### Analysis

Image analysis was performed using ImageJ^[52]^ or Fiji.^[53]^ The statistical analysis for the viability of 3D-differentiation of lt-NES, mRNA expression and survival of ejected lt-NES was performed using Origin Pro (OriginLab, USA). N designate individual hydrogel replicates, and P-values were derived using linear mixed models (LMM).

## Results and Discussion

### Hydrogel design

We developed a modular approach based on copper-free click chemistry to generate HA-based LN-521 functionalized hydrogels with tunable stiffness for neural cell encapsulation. Hyaluronan-poly(ethylene glycol) (HA:PEG) hybrid hydrogels were prepared by combining bicyclo[6.1.0]nonyne (BCN) modified HA (∼100 kDa) and an 8-arm PEG with terminal azides ((PEG-Az)_8_) as previously described.^[50,51]^ The strain-promoted azide-alkyne cycloaddition (SPAAC) reaction between BCN and azide is rapid, allows for efficient and tunable cross-linking^[29,54]^ and results in optically transparent hydrogels (Figure S1, Supporting Information). To obtain hydrogels with a modulus in the range of neural tissue (G’∼100 – 1000 Pa)^[55,56]^ we prepared the hydrogels at a ratio of BCN to azide of 10:1 and a concentration of 1 and 2 % (w/v) of the polymers (Figure 1a), which also preserved a sufficient amount of BCN groups (∼90 %) for coupling of LN. For conjugation of LN to the hydrogels, LN-521 was first modified with azide (Az) groups using carbodiimide chemistry.^[57]^ The Az groups were coupled to LN using linkers of two lengths to optimize LN conjugation and retention in the hydrogels. The longer linker comprised four ethylene glycol (EG) units, and the shorter was based on a three-carbon linker, referred to as LN-p-Az and LN-Az, respectively (Figure 1b). After purification, LN-Az/LN-p-Az (LN-(p)-Az) was combined with HA-BCN to allow for HA-BCN to bind to LN. We then cross-linked the constructs by the addition of (PEG-Az)_8_ (Figure 1a). In the absence of LN, the storage modulus (G’) of the hydrogels was approximately 350 Pa and 650 Pa for 1 % and 2 % (w/v) hydrogels, respectively, which is in the desired range for the culture of neurons (Figure 1c-d). Previous works by Saha et al.^[58]^ and Banerjee et al.^[59]^ have demonstrated that hydrogels in this stiffness range enhance proliferation and differentiation of neural stem cells (NSCs) compared to when cultured in stiffer gels. The addition of LN-Az or LN-p-Az (0.83 µM) did not have any significant effect on the stiffness of the hydrogels (Figure 1c-e) nor the gelation kinetics (Figure 1f). The gelation point (G’= G’’) was reached directly after mixing the components at 37 °C in PBS, and the hydrogels reached close to final stiffness in about 20 min. This time frame is sufficiently fast to prevent cell sedimentation while allowing enough time for handling and bioprinting. In previous work, we have also observed that the gelation kinetics for SPAAC cross-linking is highly temperature dependent and gelation can be delayed significantly when performed at room temperature and almost completely inhibited at 4°C, ^[29]^ which can further facilitate processing of the hydrogels.

**Figure 1.**
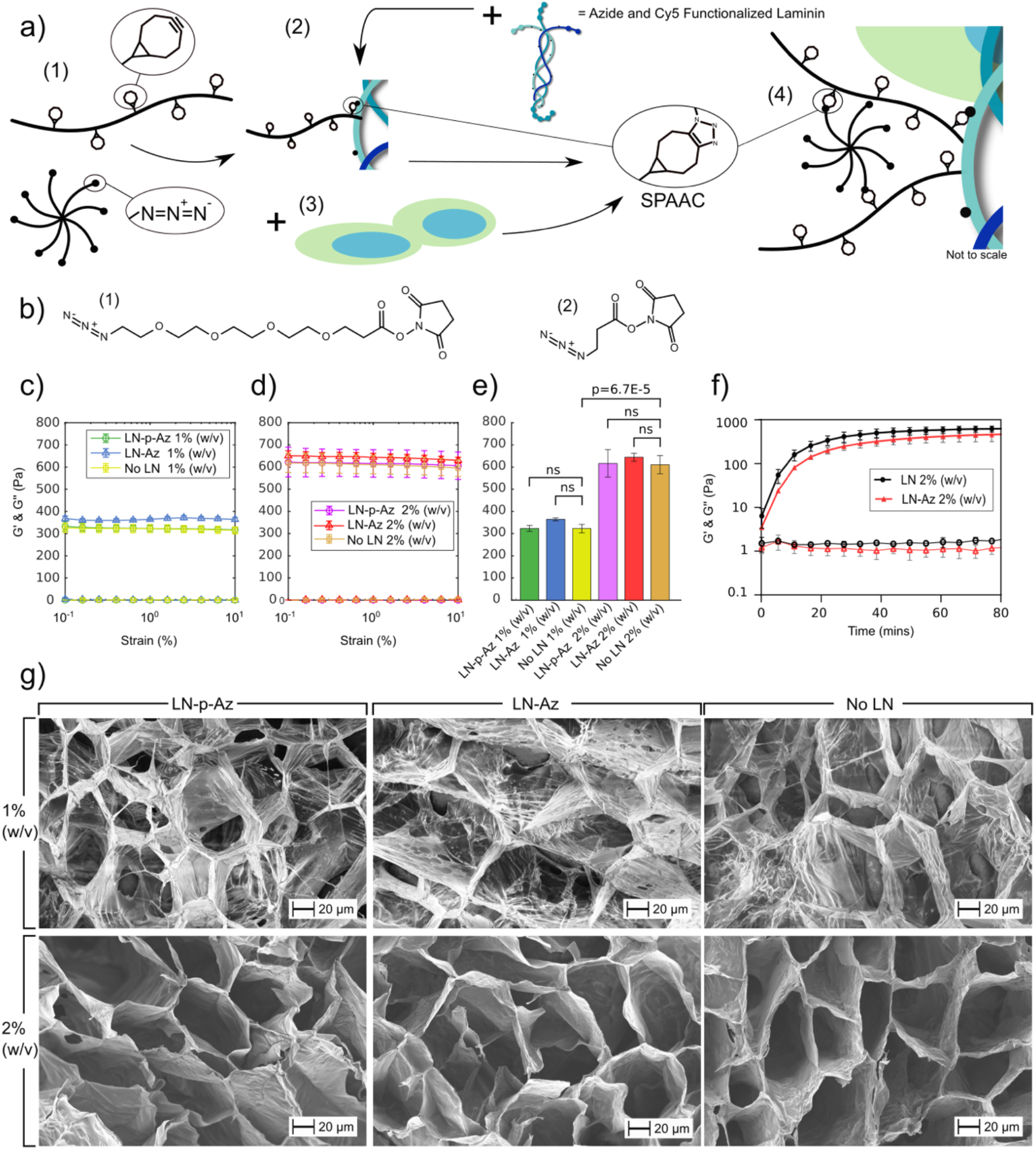
a) Schematic illustration of the modular HA:PEG-LN hydrogel system: (1) The hydrogels were crosslinked by SPAAC by combing HA-BCN and 8-armed-PEG-azide ((PEG-Az)_8_). (2) For conjugation of LN to the hydrogels, Az-functionalized LN was first conjugated to HA-BCN prior addition of (PEG-Az)_8_ to generate HA:PEG-LN. (3) For cell encapsulation, cells were combined with (PEG-Az)_8_ and then mixed with HA-BCN ± LN. LN was also labeled with Cy3 to determine conjugation efficiency and facilitate visualization of LN distribution. b) LN was functionalized with azide-terminated amine-reactive molecules with (1) a 4 EG unit linker and (2) a shorter 3 carbon linker. Oscillatory strain sweeps of HA:PEG-(LN) hydrogels with concentrations of c) 1 % (w/v) and d) 2 % (w/v). e) No significant difference in G’ at 1 % strain was seen for any of the conditions at the same hydrogel concentration. f) Hydrogel gelation kinetics. N=4 for strain sweep and gelation kinetic measurements, where each is a separate hydrogel. g) Scanning electron micrographs of hydrogels with and without LN-(p)-Az.

Scanning electron micrographs of freeze-dried hydrogels did not reveal any substantial differences between the LN and non-LN-containing hydrogels (Figure 1g). All hydrogels showed large hexagonal and interconnected pores, 50-100 µm in size. The lower weight percentage hydrogels, 1 % (w/v), showed thinner pore walls and a more fibrillar structure than hydrogels prepared at a concentration of 2 % (w/v). Pores in this size range can facilitate cell migration and cell-cell contacts and diffusion of oxygen, nutrients, and other critical factors for cell survival, proliferation, and function without a vascular system.^[60,61]^ The porous microarchitecture can also influence and promote neurite outgrowths.^[61,62]^

### Laminin distribution and retention

To characterize the influence of Az modification and linker length on the retention of LN in the hydrogels, we further labeled the LN with Cy3. Fluorescence micrographs of the hydrogels functionalized with Cy3-labeled LN-(p)-Az show a homogenous distribution of the LN with a small number of visible aggregates (Figure 2a). Based on the relative intensity of LN-Cy3 from the fluorescence micrographs, we can conclude that about twice as much LN was conjugated to the 2 % (w/v) compared to the 1 % (w/v) hydrogels and that LN-p-Az features more efficient conjugation compared to LN-Az (Figure 2b). To further determine the effectiveness of the conjugation strategies, we monitored the cumulative release of LN for 7 days (Figure 2c). For the non-Az functionalized LN, a substantial burst release was seen over the first 24 hours, corresponding to about 40 % of the incorporated LN. However, after the initial burst release, limited further release was observed, and a large fraction of the non-conjugated LN was consequently physically trapped in the hydrogel. This is likely due to the high molecular weight of LN (∼850 kDa), resulting in a slow diffusion in the hydrogel polymer network. However, LN modified with Az via both the shorter (LN-Az) and the longer and more flexible linker (LN-p-Az) was found to be substantially more efficiently retained in the hydrogels with a cumulative release of less than 5 % for the 2 % (w/v) HA:PEG hydrogels, indicating successful conjugation to the HA backbone.

**Figure 2.**
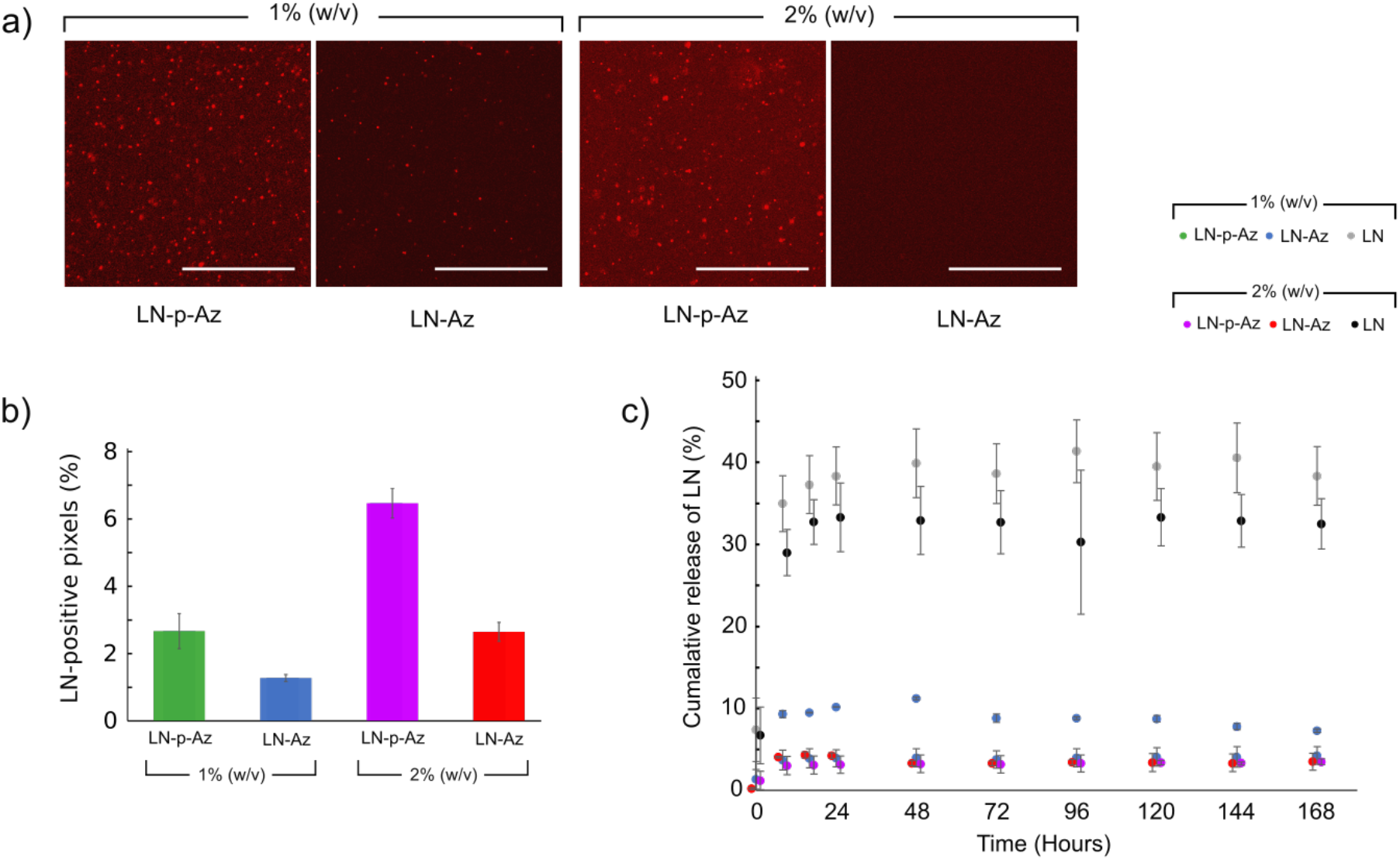
a) Fluorescence micrographs of the Cy3-labeled LN-(p)-Az conjugated in the HA:PEG hydrogels. Scale bars: 100 µm. b) LN-positive pixels determined from fluorescence micrographs of the hydrogels, N=4 for each condition. c) Cumulative release of unbound LN, LN-p-Az and LN-Az from hydrogels with 1 % and 2 % (w/v) for 7 days, N=6 for each condition, where each measurement is a separate hydrogel.

### Encapsulation and 3D culture of SH-SY5Y cells

To investigate and optimize the hydrogel system for neural cell encapsulation, we employed the human neuroblastoma cell line SH-SY5Y. This cell line has been used extensively as a model system for neurodegenerative disease in two- and three-dimensional (2D, 3D) cultures.^[63–66]^ We cultured undifferentiated SH-SY5Y cells in the hydrogels with and without LN. Samples imaged at time points of 1-, 3-, and 7-days showed even cell dispersal across all conditions with cells forming small multicellular spheroids within the hydrogels with few truncated processes (Figure S2), which is characteristic for undifferentiated SH-SY5Y cells. We used confocal imaging to confirm the spheroid-like morphology observed (Figure 3a and Figure S3, Supporting Information). The viability of the encapsulated cells was determined using an Alamar Blue (AB) assay which revealed high metabolic activity for the cells over 10 days, with no or minor differences between the conditions concerning LN-functionalization hydrogel concentration (Figure 3b). Thus, with or without the added functionalization of LN, the HA:PEG hydrogel system can efficiently sustain the neuroblastoma cells. Proliferation decreased in all conditions after day 7, which we hypothesize is a consequence of an increasing spheroid diameter over time, that might lead to oxygen and nutrient starvation of cells in the core of the spheroids, resulting in necrosis.^[67]^ The non-significant effects of LN on SH-SY5Y cell proliferation indicate that cells may adhere to the hyaluronan backbone, making any other interactions with LN redundant. The main cell surface receptor for binding to HA is CD44, a family of transmembrane cell surface glycoproteins that plays an important role in cell-cell and cell-matrix interactions and is linked to the tumorigenic properties of neuroblastoma cells. Subpopulations of SH-SY5Y cells have been shown to express CD44.^[68]^ As indicated by immunostaining, the undifferentiated SH-SY5Y cells express CD44 when cultured in HA:PEG, both in the absence and presence of LN-p-Az, providing additional means for cell adhesion to the hydrogels in addition to integrin-LN interactions (Figure S4, Supporting Information).

**Figure 3.**
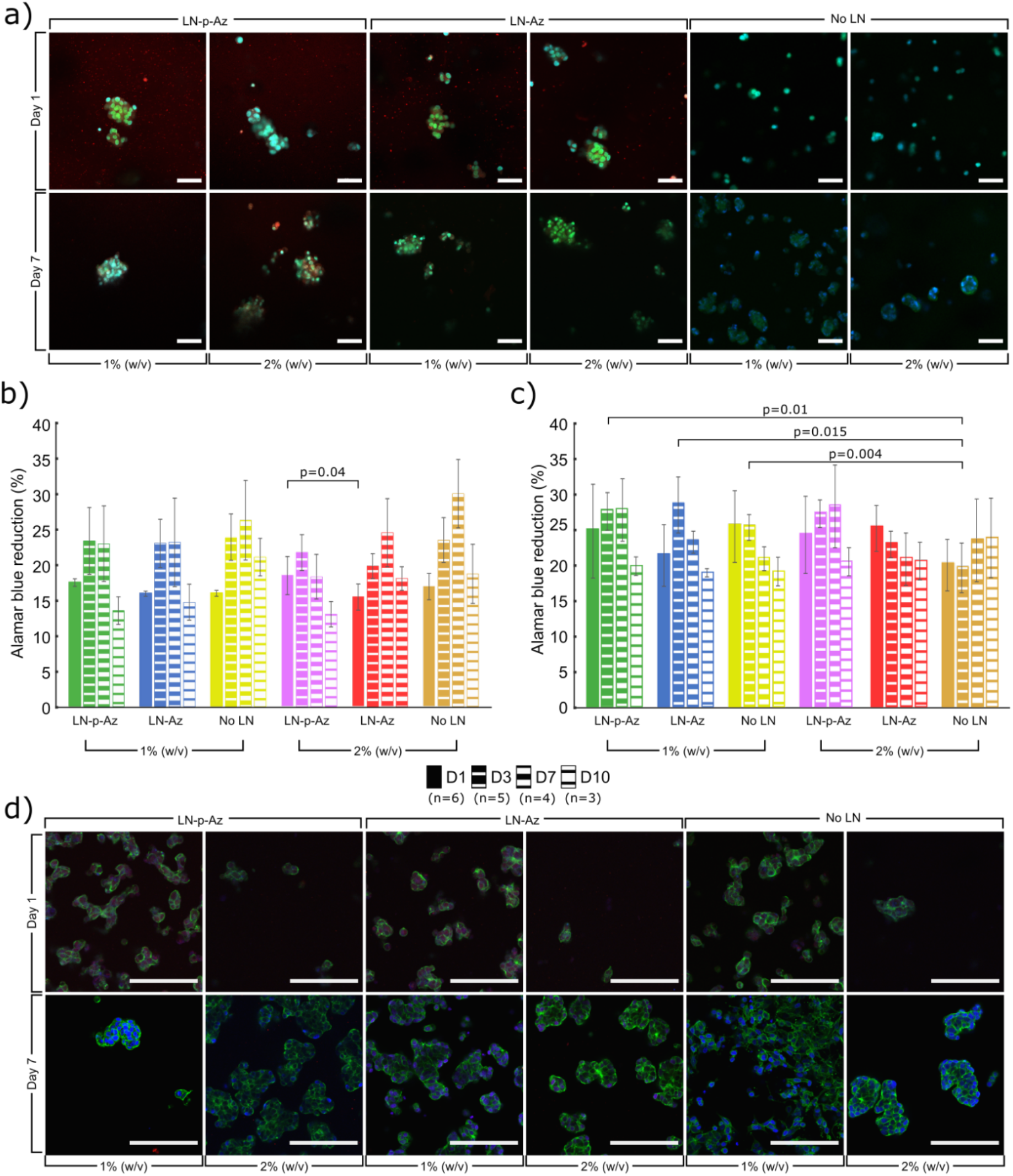
a) Confocal images of undifferentiated SH-SY5Y cells cultured in 1 and 2 % (w/v) HA-BCN:PEG hydrogels with and without LN-(p)-Az, stained for F-actin (Phalloidin, green) and Hoechst nuclear dye (blue). Also shown, Cy3 fluorescence labeling of laminin (red). Alamar blue cell viability of (b) undifferentiated SH-SY5Y and (c) RA treated differentiated SH-SY5Y cells, cultured in HA:PEG hydrogels with and without LN-p-Az, N= as indicated on figure legend for each condition, where each is a separate hydrogel. d) Confocal images of RA treated differentiated SH-SY5Y cells cultured in 1 and 2 % (w/v) HA:PEG hydrogels with and without LN-p-Az, stained for F-actin (Phalloidin, green) and Hoechst nuclear dye (blue). Scale bars: 100 µm. Significance was calculated using one-way ANOVA Tukey HDV.

To induce differentiation of the SH-SY5Y cells, retinoic acid (RA, 10 µm) was added to the cell culture medium 7 days before encapsulation in the hydrogels. RA has highly potent growth-inhibiting and cellular differentiation-promoting properties and triggers differentiation, primarily to a cholinergic neuronal phenotype.^[69,70]^ The differentiated cells remained viable in all hydrogels up to 10 days in culture as indicated by the AB assay (Figure 3c). Interestingly, the viability of the differentiated SH-SY5Y cells rapidly decreased in hydrogels supplemented with non-conjugated LN (Figure S5, Supporting Information). RA differentiation of SH-SY5Y cells increases the expression of α_3_β_1_ integrin heterodimers,^[71,72]^ which interact strongly with LN-521.^[73,74]^ Binding of non-conjugated LN-521 thus likely interferes with cell-hydrogel binding, triggering cell death via anoikis pathways.^[75]^ Confocal images of the differentiated cells showed homogenously distributed cell clusters in all conditions, growing into larger spheroids by day 7 (Figure 3d), similar in size and geometry as SH-SY5Y cultured hydrogels of collagen or alginate.^[76,77]^

### lt-NES viability during spontaneous 3D differentiation

Whereas the 3D cultures of neuroblastoma cells represent an important neural disease model, the potential to encapsulate and culture lt-NES in the HA-PEG hydrogels offers opportunities also to explore models of healthy tissues, advanced models of genetic disorders, and development of cell-based therapeutic strategies. Neural progenitor cells and lt-NES spontaneously differentiate into mixed cultures of high (80-95 %) percentage neurons and some glial cells. lt-NES have been used in several studies both as disease models and as a source to create healthy neurons, both in 2D and 3D.^[78–80]^ Here, lt-NES were encapsulated in the HA:PEG hydrogels with the addition of LN, with and without an Az-conjugation. To benchmark these defined hydrogels as a matrix for cultivation and differentiation of the sensitive lt-NES (pre-differentiated in 2D for 5 days), we used commercially available Matrigel. After 1 day of the subculture in the hydrogels, we measured lt-NES metabolic activity using AB, demonstrating viable cells in all gel conditions (Figure 4a). Whereas we observe no or only minor differences in viability of the conditions with/without LNs, the viability is about 2-3 times higher in Matrigel. Matrigel is tumor derived, composed by several partially fragmented ECM proteins, and contains traces of growth factors that can give higher cell survival and proliferation of the lt-NES throughout the first 24 h of culture as confirmed by brightfield microscopy (Figure S8, Supporting Information).

**Figure 4.**
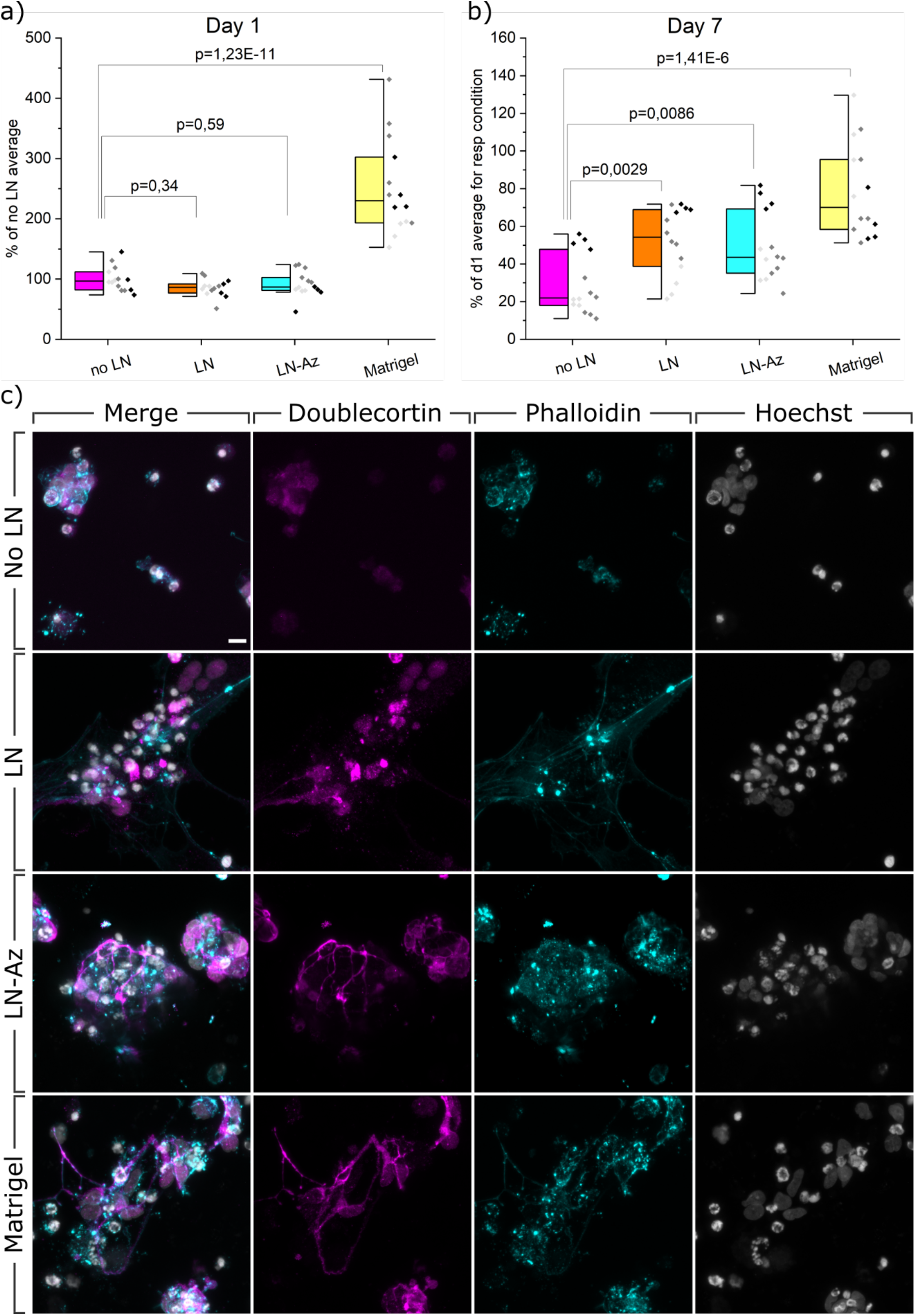
AB viability assay and confocal micrographs of spontaneous 3D-differentiation. a) Viability measured with Alamar blue after 1 day of differentiation of lt-NES in respective hydrogels. b) Viability measured with Alamar blue after 7 days of differentiation of lt-NES in respective hydrogels. Data was collected from three individual experiments (indicated with different shades of grey), N=14, where N represents one hydrogel replicate. p-values were derived using LMM with all data included. c) Confocal micrographs of spontaneous 3D-differentiation in respective hydrogels. lt-NES are stained with neuronal marker Doublecortin (magenta), cytoskeletal f-actin marker Phalloidin (cyan) and nuclear stain Hoechst (grey). Scale bar 10 µm.

Additionally, we observe that the Matrigel encapsulated cells result in a larger spread of data points in viability compared to the other hydrogel conditions with an IQR of [193;302], compared to hydrogels without LN [82;112], non-conjugated LN [77;112] and LN-Az [81;103]. The well-defined HA:PEG hydrogels thus clearly provide better reproducibility between independent experiments than Matrigel (Figure 4a). After 7 days of subculture in the hydrogels (Figure 4b), a significant difference in viability was observed in hydrogels containing LN. However, in contrast to the differentiated SH-SY5Y cells, conjugation of the LN does not appear to change the viability of the cells compared to non-conjugated LN. Similar to what we observe after 1 day of subculture, the viability of the cells in Matrigel is significantly higher and shows a larger spread of the individual data points (seen as different shades of grey). An analysis of the IQR shows that Matrigel had the largest variability [58;96] compared to the conditions with no LN [35;69], LN [39;69], and LN-Az [18;48]. We hypothesize that the increase in viability in Matrigel compared to day 1 is partly due to a continuous proliferation throughout the 7 days of subculture, in line with the higher proliferative potential of that material as mentioned above, caused by its various components such as mixed ECM proteins and growth factors traces.

### Neural Stem Cell morphology after 3D differentiation

More specific assessments of neuronal differentiation require characterization of neuronal markers both on an imaging and mRNA expression analysis level. As seen by immunocytochemistry in (Figure 4c), lt-NES cultured in the HA:PEG hydrogels without LN are generally characterized by singularly distributed cells and smaller clusters of cells. The expression of Doublecortin (DCX), an early marker of neuronal differentiation, is mainly limited to the area surrounding the individual nuclei, and occasional neurite outgrowths are found. Phalloidin (F-actin stain) shows how the soma is rounded up, and little cell spreading or contact with the hydrogel is seen. Singularized nuclei appear brighter and condensed to a higher level compared to clustered nuclei. A larger proportion of loosely clustered single cells is observed when adding LN to the hydrogels, while some smaller clusters are still present. Similar to the no LN condition, DCX expression is limited to an area around the cell nuclei. The cells form small clusters with connected neurites, confirmed by F-actin staining (Phalloidin). The F-actin staining further visualizes cell-cell connections and close interaction between the cells and the-hydrogel. In the LN conditions mixed morphologies of nuclei are observed, some are larger and more oval-shaped, and others are brighter and more condensed close to pycnotic, much like those seen in the condition no LN. In the HA:PEG hydrogels with conjugated laminin, Az-LN, the cell distribution appears similar to the LN condition, with respect to singular cells and small clusters. DCX expression apart from the soma is detectable in neurite outgrowths connecting cells in the clusters. F-actin staining reveals a somewhat condensed cytoskeletal structure indicating strong cell-cell interactions. Most nuclei appear larger and oval-shaped when in the loose clusters, indicating that cells are mostly healthy, although some nuclei are brighter and more condensed. In the Matrigel condition, cells are growing both singularized and in less tightly packed clusters, and we observe that the cells send out longer processes between each other. The expression of DCX extends from the soma to the processes that form a network-like structure. The spreading and outreach of the cells are confirmed by the cytoskeletal constructs seen in Phalloidin, suggesting that the cells in a similar way to the LN condition can attach to their microenvironment. Our findings support previous data showing that Matrigel can allow the formation of complex neuronal networks.^[81]^

Depending on the size and format of the 3D-hydrogel being stained, lengthy immunostaining protocols are needed, compared to 2D cultures, to provide enough diffusion time for antibodies to penetrate the gel and bind to the cells. With Matrigel, we found it difficult to avoid unspecific binding and high background noise even with repeated and longer washing steps. Issues like these can prove disruptive to imaging, especially image analysis, since larger clusters of background noise can be easily similar in size to thin neurites or other structures of interest in neural stem cell cultures. Notably, we did not experience background noise issues and unspecific binding with HA:PEG-based hydrogels, which, on top of its defined formula, gives HA:PEG one more advantage over Matrigel.

### mRNA expression analysis

After 5 days of spontaneous pre-differentiation in conventional cell culture flasks and 7 days of continued spontaneous differentiation in 3D-hydrogels, we extracted RNA in the typical range of 5-20 ng/µl from 50 µl hydrogels. We observe significant changes in gene expression in most of the analyzed genes, both between some hydrogel conditions (Figure 5) and compared to the undifferentiated state of lt-NES (Figure S6, Supporting Information). The neuronal marker DCX (Figure 5a) is upregulated (1.75-2.75 times) in all hydrogel conditions and more so when LN was added to the hydrogels. However, we see no significant effect of LN conjugation. Cells differentiated in Matrigel showed the highest upregulation of DCX, supported by our imaging findings (Figure 4c). When compared to the undifferentiated state of lt-NES, we observe that all conditions had significant upregulation of DCX (Figure S6, Supporting Information), which is in line with previous studies where neuroepithelial stem cells show expression of DCX after 7 days of neuronal differentiation.^[81]^ The later neuronal marker Tubulin Beta 3 Class III (TUBB3) shows a slight upregulation (1-1.5 times) compared to undifferentiated lt-NES (Figure S6, Supporting Information), where Matrigel again had the highest level of upregulation. However, when comparing expression between the different hydrogel conditions (Figure 5b), we do not see any significant up or downregulation when adding LN or LN-az. We observe the only significant change in Matrigel, where TUBB (1.25-2 times) was slightly upregulated. As a measure of cell attachment and interaction with the hydrogels, including synaptogenesis, we investigate the expression of Synapsin 1 (SYN1). We observed an upregulation in all conditions where the addition of LN seems to make a slight difference in upregulation but is not affected by the Az conjugation (Figure 5c). The highest upregulation we see in Matrigel (1.75-3.25 times).

**Figure 5.**
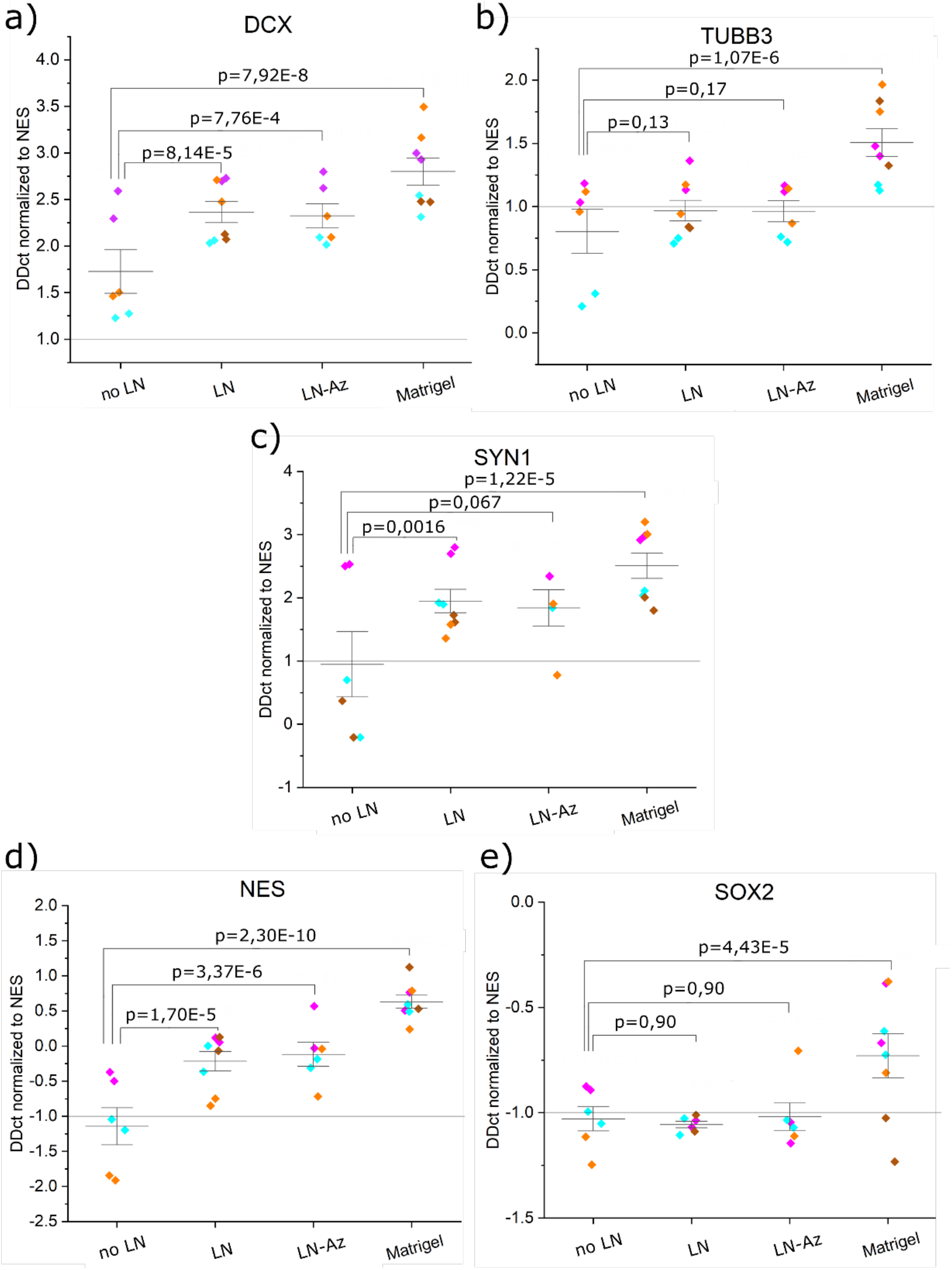
mRNA analysis profile of spontaneous 3D-differentiation of lt-NES in different hydrogel conditions. a) DCX, b) TUBB3, c) SYN1, d) NES, and e) SOX2. All samples contain the housekeeping gene GAPDH. P-values were derived with LMM, and data were normalized to HA:PEG. Data is collected as duplicates from three or four independent experiments (indicated by different colors).

As another measure of neuronal differentiation, we include two stem cell markers, Nestin (NES) and SRY-Box Transcription Factor 2 (SOX2), expecting that these two genes should be downregulated in the case of successful neuronal differentiation. SOX2 has a critical role in maintaining pluripotency and directing pluripotent stem cells to neural progenitors.^[82]^ A previous study using human neuroepithelial stem cells to create human midbrain organoids reported a change from 35 % to 18 % positive SOX2 cells when comparing expression at 27 days and 61 days of differentiation, respectively.^[83]^ We have seen in prior work that spontaneous differentiation of lt-NES in 2D-culutre result in downregulation of SOX2 after 28 days.^[84]^ Our results show significant downregulation of NES in the HA:PEG hydrogels without LN, but not when any type of LN was added. Compared to the no LN condition, we observe a slight upregulation of NES in the Matrigel condition, indicating that the stem cell state would be more preserved for the cells cultured in Matrigel. As for SOX2, we see a clear downregulation in all hydrogel conditions, with no difference if any kind of LN is added. The data has high variability in the Matrigel condition, and no significant downregulation can be concluded in terms of change in DDct values. Similar to our observations in SOX2 regulation, such high variability and lack of downregulation of NES, as we see in the other hydrogel conditions, gives Matrigel a disadvantage as a matrix for neuronal differentiation compared to the HA:PEG-based hydrogels.

For all the genes, we observe the largest variation in data points from the no LN condition, which was also the condition that, in general, had the lowest mRNA yield.

### Ejection of NES in HA:PEG and media

In addition to providing an excellent matrix for 3D culture and neuronal differentiation, the defined composition combined with the bioorthogonal cross-linking chemistry of the HA:PEG hydrogel can facilitate implementation of cell-based regenerative therapies.^[85–88]^ Syringe-based cell transplantation exposes the cells to significant shear forces that may mechanically disrupt the cells and substantially reduce cell viability. In many transplantation studies, PBS or cell media is used as a vehicle to carry the cells. However, shear-thinning hydrogels have been demonstrated to provide a protective effect during the injection.^[89]^ To investigate the ability of the HA:PEG hydrogels to protect cells experiencing sheer force when ejected through a syringe needle, we compared the viability of lt-NES in an HA:PEG matrix to cell media and a collagen gel both in an acute state and after 24 h by assessing the amount of live and dead cells. At the acute state, we observed reduced viability of the HA:PEG ejected cells compared to those ejected in cell media. The same effect was seen when comparing cells ejected in a Collagen Matrix compared to cell media (Figure 6, Figure S7 Supporting Information). However, after 24 h, cells ejected in the HA:PEG matrix showed higher viability than cells ejected in cell media, suggesting that the hydrogel provides a protective environment during and after syringe ejection.

**Figure 6.**
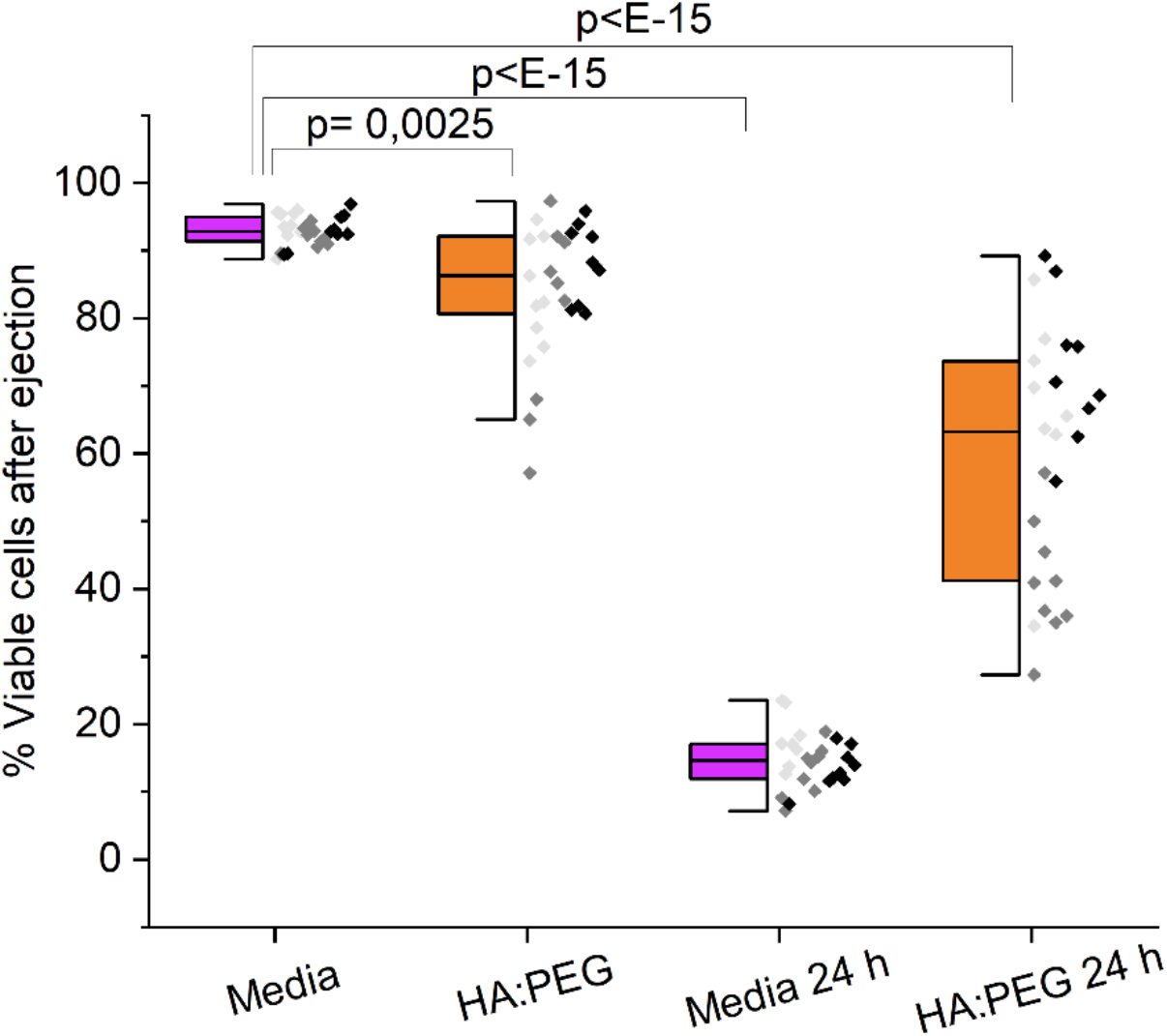
Survival of ejected lt-NES in HA:PEG and cell media immediately after ejection and after 24 h measured with LIVE-DEAD assay. Data was collected from three individual experiments (indicated with different shades of grey), N=27, where N represents one replicate of ejected cells. Outliers were removed with Grubbs Test, and p-values were derived using LMM in Origin Pro.

### 3D Bioprinting

The protective effect of the HA:PEG hydrogels on cells during syringe ejection is also a highly attractive feature for 3D bioprinting applications. To assess the printability of the hydrogels, hydrogel lattices (1 × 1 cm) were fabricated using a Cellink BioX bioprinter (Figure 7a). In addition to enabling printing of features with dimensions < 400 µm, the hydrogels supported the high viability of the bioprinted SH-SY5Y cells (Figure 7b,c). Similar to syringe-based cell ejection, 3D bioprinting exposes cells to substantial shear forces that can be detrimental for cell viability due to the rapid change in fluid velocity when the cell suspension is forced from the syringe into the much smaller diameter needle, resulting in cell rupture.^[90,91]^ By encapsulating the cells in the HA:PEG hydrogels matrix, the cells were protected from the lethal shear forces during bioprinting. SH-SY5Y cells encapsulated in LN-functionalized HA:PEG showed high (> 85 %) viability 24 h after printing, similar to cells carefully extruded through a pipette (Figure 7c) and on par with carefully optimized alginate-based bioinks.^[92]^ Moreover, the bioprinted SH-SY5Y cells showed a similar morphology and distribution in the 3D bioprinted structures as when cultured in the casted hydrogels (Figure 7b), indicating the potential of this hydrogel system for 3D bioprinting of neural disease models. Interestingly, 3D bioprinting of the SH-SY5Y cells in HA:PEG hydrogels functionalized with LN resulted in more spheroids than hydrogels without LN (Figure 7d).

**Figure 7.**
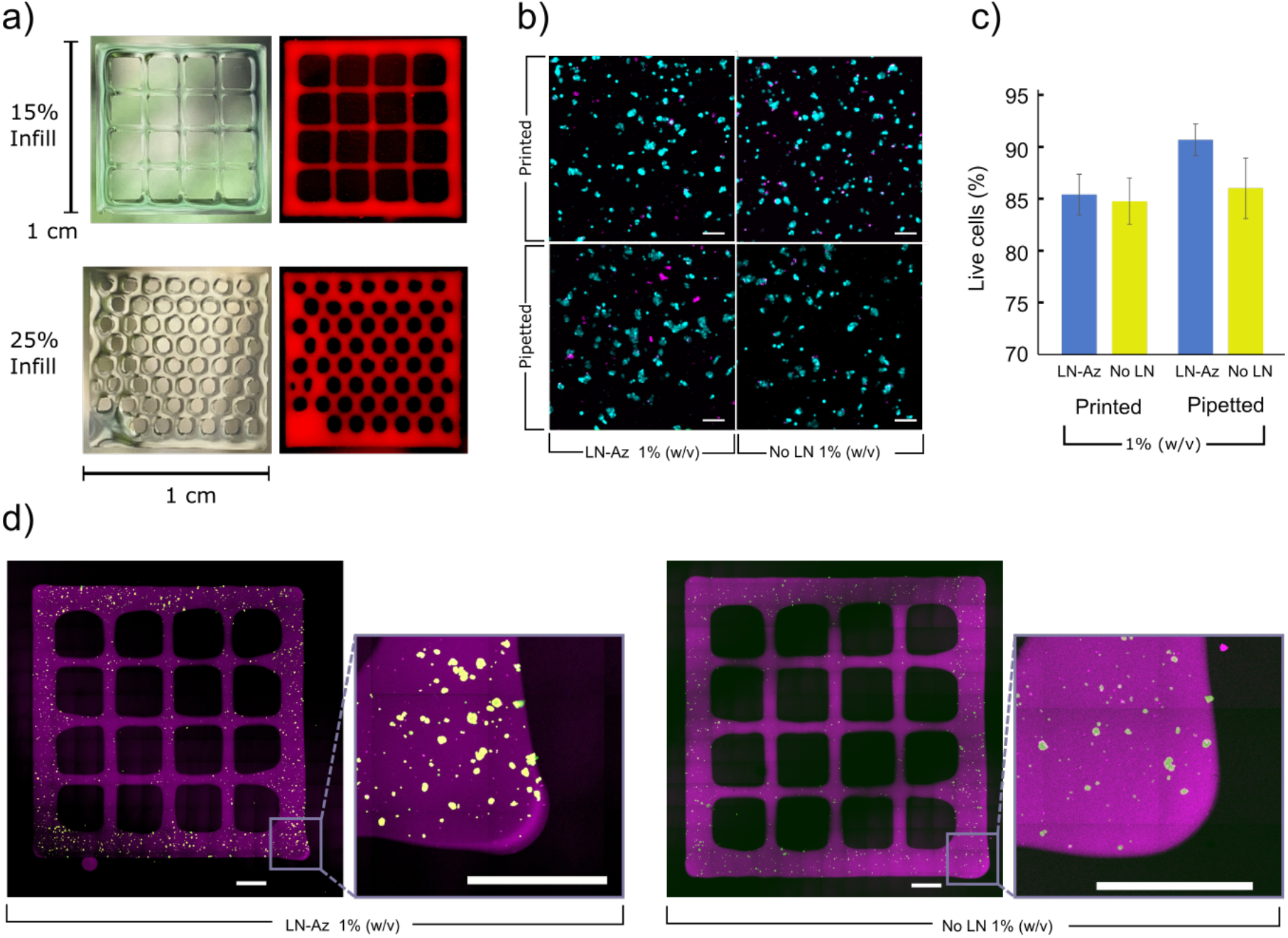
a) 3D bioprinted structures based on the HA:PEG-LN hydrogels at a concentration of 1 % (w/v). Hydrogels (red) were dyed with Cy5 and illuminated using a white light source. b) Live (Cyan)/dead (Magenta) staining of SH-SY5Y cells 24 h after bioprinting. c) SH-SY5Y cell viability 24 h after bioprinting or pipetting when encapsulated in either HA:PEG (without LN) and HA:PEG-LN hydrogels at a concentration of 1 % (w/v), N=4 for each condition. d) SH-SY5Y cell bioprinted into grid structures (purple) of Cy5-labeled HA:PEG-LN and HA:PEG, respectively, at a hydrogel concentration of 1 % (w/v) and imaged using tiled confocal microscopy 24 hours after bioprinting. Encapsulated SH-SY5Y were stained using Live/Dead staining Live (Cyan)/dead (Magenta). Inset square indicates a magnified portion. Scale bars are 100 µm.

## Conclusions

We presented a tunable and modular HA-based LN-521 functionalized hydrogel that can effectively retain LN over 7 days, showing a successful conjugation of the LN to the hydrogel backbone. We demonstrated that the hydrogel supports the growth and differentiation of the widely used neural cell model SH-SY5Y. The SH-SY5Y showed high viability after 10 days of subculture in the hydrogels and appeared to grow in clusters according to actin staining and immunostaining of the HA-receptor CD44. We further demonstrated that more sensitive and advanced cell model lt-NES successfully can be spontaneously differentiated to neuronal fates and develop neurite outgrowths in these hydrogels. According to viability assays, LN does not support the cells’ immediate (24 h) survival but does change the viability on a long-term culture of 7 days. Our mRNA expression analysis suggests a slight upregulation in neuronal markers DCX, TUBB3 and SYN1 with the addition of LN to the hydrogels. Our data also suggests that stem cell marker SOX2 is slightly downregulated, whereas we see no significant difference in the expression of the stem cell marker NES with the addition of LN. We proved the possibility of ejecting lt-NES through a 27G syringe and that adding HA:PEG as an ejection matrix will protect the cells by higher survival after 24h, compared to cells ejected in cell media. This protective effect of the hydrogel matrix could not be measured through viability at an immediate stage. We furthermore successfully bioprinted SH-SY5Y cells encapsulated in LN-functionalized HA:PEG with >85 % survival after 24 h. We also observed that the bioprinted cells maintained the same morphology as when cultured in 3D gels, and surprisingly we found that conjugating LN in the hydrogels promoted the formation of spheroids to a larger extent than without the added LN after bioprinting.

In summary, we presented a defined, bioprintable, tunable hydrogel system allowing controlled covalent conjugation of the full-size essential ECM molecule laminin. The hydrogel is compatible with sensitive neural stem cells that could be used in advanced tissue and disease models of the developing brain and offer higher reproducibility and simplifies imaging, compared to conventional biologically derived hydrogel systems. Possibilities to process the materials and protect cells during syringe ejection can further facilitate the development of advanced neuronal tissue and disease models and clinical translation of neuronal cell therapies.

## Supporting information

Supporting data and detailed experimental section

## Acknowledgments

M. Jury and I. Matthiesen contributed equally to this work. BioLamina AB kindly provided BioLaminin 521. The financial support from the Swedish Foundation for Strategic Research (SFF) (FFL15-0026), the Swedish Government Strategic Research Area in Materials Science on Functional Materials at Linköping University (Faculty Grant SFO-Mat-LiU No. 2009-00971), the Carl Tryggers Foundation, and the Knut and Alice Wallenberg Foundation (KAW 2016.0231 and 2020.0206) is gratefully acknowledged.

