## Supporting data and detailed experimental section for "Bioorthogonally Cross-Linked Hyaluronan-Laminin Hydrogels for 3D Neuronal Cell Culture and Biofabrication"

### Materials and Methods

**General:** Hyaluronic acid (150 kDa) was obtained from Lifecore Biomedical (Minnesota, USA). Human recombinant laminin, Biolaminin 521 (LN-521), hereafter termed LN, was obtained from BioLamina AB (Stockholm, Sweden). Unless otherwise noted and used without further purification, all other chemicals were purchased from Sigma Aldrich (Missouri, USA). All cell culture reagents were purchased from Thermo Fisher, USA, unless stated otherwise.

**Laminin labeling:** LN was modified with azide moieties (Az) using either azido-dPEG$_8$-NHS (long linker) or a short amine-reactive azide-functionalized linker. The latter was obtained by reacting azidopropionic acid (0.17 mmol) with N-hydroxysuccinimide (NHS, 0.17 mmol), 1-Ethyl-3-(3-dimethylaminopropyl)carbodiimide (EDC, 0.17 mmol) dissolved in 2 ml chloroform with 100 µl of N, N-Diisopropylethylamine (DIPEA). After one hour, the solvent was evaporated, and the remaining powder was suspended in 230 µl of DMSO to give a solution of 375 mM. Next, LN was run through a G10 mini trap column (Cytiva Life Sciences, USA) to remove tris buffer, replacing it with 100 mM $NaHCO_3$ pH 7.9 before conjugating the azide-linkers. To label the LN with Az and fluorophore, 10 µl of 375 mM of the linkers and 0.75 µl Cy3-sulfo-NHS (DMSO, 5 mM) was added to 1 ml LN (1.25 µM) and incubated for 30 min. Next, Tris buffer (10 mM, pH 8, 10 µl) was added to quench the remaining NHS. Finally, the modified LN was purified on a PD10 midi trap column (Cytiva Life Sciences, USA), and the collected product was suspended in 1.5 ml phosphate-buffered saline (PBS, 137 mM NaCl, 2.7 mM KCl, 8 mM $Na_2HPO_4$, and 2 mM $KH_2PO_4$) to give a final concentration of 0.83 µM. The As-functionalized LN is hereafter defined as LN-Az for the short azidopropionic acid-derived linker and LN-p-Az for the long azido-dPEG$_8$-NHS ester derived linker.

**Synthesis of HA-BCN:** HA was modified with bicyclo[6.1.0]nonyne (BCN) as described in detail in Selegård et al. [1]. Briefly, N-[(1 R,8 S,9 S)-Bicyclo[6.1.0]non-4- yn-9-ylmethyloxycarbonyl]-1,8diamino-3,6-dioxaoctane (BCN-$NH_2$) was dissolved in 5:1 (v/v) acetonitrile:MilliQ water (18.2 Ω $cm^{-1}$) before the addition of EDC and 1-hydroxybenzotriazole hydrate. This was combined with HA dissolved in 2-(N-morpholino)ethanesulfonic acid (MES) buffer (100 mM, pH 7) and the carbodiimide reaction was allowed to continue for 24 h, after which time dialysis (MWCO 12–14 kDa, Spectra/ Por RC, Spectrum Laboratories Inc.) was carried out in acetonitrile: MilliQ water (1:10 v/v) for 24 h and then MilliQ water for three days. The dialyzed product, referred to as HA-BCN, was lyophilized and stored at −20°C. HA-BCN had a derivatization degree of approximately 19 % based on $^{1}$H-NMR analysis.

**Hydrogel formation:** HA-BCN was suspended in PBS or appropriate cell culture media to give a concentration of 1 % (w/v) or 2 % (w/v). LN-Az or LN-p-Az was added in the proportion of 10 % of total hydrogel volume, thoroughly mixed and sealed in a microtube and incubated at 37°C for one hour. After this time, an 8-arm Az-terminated polyglycol ethylene crosslinker ($(PEG\text{-}Az)_8$) was suspended in PBS at the same concentration as HA-BCN and added to the LN-functionalized HA-BCN at a ratio of 0.724:0.276 HA-BCN:$(PEG\text{-}Az)_8$. The mixture, including cells where relevant, was thoroughly but quickly mixed, and the gels were formed as required. In the case of the No LN condition, PBS was used in place of LN.

**Rheology:** Oscillatory rheology was conducted using a Discovery HR-2 rheometer (TA Instruments, USA). Hydrogels were first cross-linked for 1 hour at 37 °C, followed by the addition of enough PBS to cover the hydrogel and then incubated for a further 24 hours at 37 °C. The hydrogels were examined using frequency sweeps (1 % strain, 0.1 to 10 Hz) and amplitude sweeps (1 Hz, 0.1 to 10 % strain) using an 8 mm plate geometry. Gelation kinetics was investigated using a 20 mm 1° geometry at 1 Hz and 1 % strain and recorded every 6 seconds. Samples were prepared in situ on the rheometer by premixing the HA-BCN and $(Peg\text{-}Az)_8$ and pipetting 50 µl atop the loading platter, precooled to 4 °C. The geometry was lowered to operating height, the platter rapidly raised in temperature to 37 °C, and the measurements started. It is approximated that the measurements were started 10 seconds after mixing the components.

**Scanning Electron Microscopy (SEM):** Hydrogels were first cross-linked for 1 hour at 37 °C, followed by the addition of enough phosphate buffer (PB, 10 mM, pH 7.4) to cover the hydrogel, and then incubated for a further 24 hours at 37 °C. The resulting hydrogels were placed onto carbon disc sample holders, rapidly frozen using liquid nitrogen and lyophilized. The dried samples were carefully sliced to expose the hydrogel structure. Next, the samples were sputter-coated with platinum for 10 seconds. SEM-analysis was conducted with LEO 1550 Gemini from Zeiss operated at a voltage of 3 kV.

**Laminin Retention:** Hydrogels were formed at a volume of 50 µl in a 96 well plate with PBS as the hydrogel component suspension buffer, with 200 µl of PBS covering the formed hydrogels. After adding the PBS and a gentle shaking, 125 µl of the buffer was removed to a 96-well clear-bottom plate, and fluorescence intensity of released Cy3-labeled LN was measured using a TECAN infinite M1000 Pro plate reader (Tecan, Switzerland) at 569 nm using an excitation wavelength of 555 nm (bandwidth of +/- 5nm). The sample was then returned to the corresponding hydrogels, sealed with parafilm and wrapped in foil. The plate was placed on a rotary plate set to 100 rpm and incubated at room temperature. The measurement procedure was repeated for each time point.

**SH-SY5Y cell culture and differentiation:** SH-SY5Y cells were obtained from ATCC. The cells were thawed and cultured in DMEM/F12 cell media (Biowest, USA) containing 10 % FBS (Biowest, USA), 1 % PenStrep (Biowest, USA) and 1 % non-essential amino acid supplements (Biowest, USA), hereafter termed SH-SY5Y maintenance media. Cells were seeded in flasks and split 1:4 when 80 % confluence was reached. Passage number was never allowed to exceed 20 above the passage number at which supplied. Differentiation of SH-SY5Y cells was performed as cells were seeded into a T-75 flask following a 1:4 split. Retinoic acid (RA) was added to maintenance media to give a final concentration of 10 µM (SH-SY5Y differentiation media). Cell media was aspirated after three days, the cells were washed with PBS, and fresh cell differentiation media was added. After seven days, the cells were considered to have undergone differentiation and were used as required.

**Viability of SH-SY5Y cells:** Hydrogels were prepared as described earlier, with SH-SY5Y maintenance media as the carrier solution. The SH-SY5Y cell culture was trypsinized (Trypsin 0.25 % in PBS, Biowest, USA) for 2 min at 37°C to ensure a homogeneous single-cell suspension and counted using trypan blue and a Bürker chamber and the appropriate number of cells were pelletized at 120g for 6 min. The $(Peg\text{-}Az)_8$ solution was added, and the cells were resuspended. The solutions of HA-BCN + LN-(p)-Az and $(Peg\text{-}Az)_8$ and cells were combined, and 50 µl hydrogels with $10^6$ cells per hydrogel were produced in 96 well plates. The plate was sealed with parafilm and incubated for one hour at 37 °C, after which 100 µl of maintenance media was added per well. The plate was placed in the incubator (37 °C, 5 % $CO_2$). All cell media and PBS used were warmed to 37°C before use unless stated. After 24 hours, the cell media was removed, the gels were washed with PBS followed by the addition of 10 % AlamarBlue™ Cell Viability Reagent (AB) (Invitrogen, USA) in 100 µl maintenance media. After one hour of incubation, the AB solution was removed to a 96 well plate and absorbance at 570 nm, and 600 nm (+/-5nm) was collected using a TECAN infinite M1000 Pro (Tecan,

Switzerland). The hydrogels were washed with PBS, and 100 µl of maintenance media was added, and the plate was placed in the incubator (37 °C, 5 % $CO_2$) until the next time point (3, 7 and 10 days). The percentage of AB reduction was calculated using the collected AB absorbance values of the hydrogel samples and unreduced AB in cell media as set out in the official protocol.

**Maintenance lt-NES:** Neuroepithelial stem cells (lt-NES)[2] C9 were provided by the iPS Core facility (Karolinska Institute) previously derived from hiPSC C9[3]. The cells were maintained in a PLO (Sigma Aldrich, USA) and L2020 (Sigma Aldrich, USA) pre-coated T25 culture flask, both coatings incubated for a minimum of 2 h, 1:500 in PBS. The cells were cultured in DMEM/F12+Glutamax supplemented with 1 % N2 0.1 % B27 supplement, ten ng/ml bFGF (R&D Systems, USA) and 10 ng/ml hEGF (Sigma Aldrich, USA), hereafter termed lt-NES maintenance media. When the cells reached a confluency of 80 %, they were passages by a 4 min TrypLE incubation at 37 °C, after which they were collected in defined trypsin inhibitor for centrifugation and replating in media at a density of 40 000 cells/$cm^2$. All cell cultures described were kept in an incubator at 37 °C with 5 % $CO_2$ and 90 % humidity.

**Spontaneous 2D-differentiation of lt-NES:** On day −1, lt-NES were seeded out in maintenance media at a density of 40 000 cells/ $cm^2$ in LN521 (BioLamina AB, Stockholm) pre-coated T75 culture flasks, incubated for a minimum of 2 h, 1:500 in PBS at 37°C. After 24 h, media was changed to DMEM/F12 supplemented with 1 % N2 and 0.1 % B27 supplement (hereafter termed lt-NES differentiation media) to induce differentiation. Half of the media was changed every 48 h. On day 5, the pre-differentiated cells were collected in DMEM/F12 with 10 % fetal bovine serum and 10 % dimethyl sulfoxide and placed in a Mr. Frosty for 24 h in −80 °C, to then be stored in liquid nitrogen until further experiments.

**Encapsulation and spontaneous 3D-differentiation of lt-NES:** As a first step, lyophilized HA-BCN and PEG-Az were reconstituted in DMEM/F12 to 2 % (w/v) for 1-2 h or until fully dissolved at room temperature. LN or LN-Az, prepared as above, or media (for control gels), was incubated with HA-BCN at 37 °C for 30 min in the proportion of 5 % of the final hydrogel volume components. The 5-day pre-differentiated cells, described in the section above, were thawed in differentiation media with 0.1 % (w/v) Y27632 (RnD Systems, USA). The cells were then mixed with the pre-incubated HA-BCN-LN at a 5 million cells/ml concentration at a ratio resulting in a final 1 % (w/v) of the hydrogel components. Next, PEG-Az was added to the HA-

BCN-LN cell mixture at a ratio of 0.724:0.276 HA-BCN:$(PEG\text{-}Az)_8$ and carefully pipetted into a half area 96 wp (Corning, USA) at 50 µl gels/well. As a comparison of the matrix, growth factor-reduced Matrigel (Corning, USA ) gels were prepared by mixing Matrigel 1:1 with cells suspended in media at a concentration of 5 million cells/ml, resulting in a final concentration of 4.3 mg/ml Matrigel. The gel-cell mixture was then carefully pipetted in the half area 96wp. The gels were left to cross-link for 1h at 37 °C. Finally, 100 µl of lt-NES differentiation media was added on top of the formed hydrogels, and the gels were kept in the incubator until further manipulation. Half of the media was changed every 24 h.

**Viability assay after spontaneous 3D differentiation:** After 1, 3 and 7 days of culture, cell survival and proliferation were measured by incubating the cells in their media with 10 % AlamarBlue$^{TM}$ for 3h at 37 °C. After the incubation, media was collected in a half area 96 wp and the fluorescence signal was measured at ex/em 550 nm/590nm, bandwidth ex/em 9 nm/ 20 nm, using a Tecan Infinite 200 Pro Multimode Plate Reader (Tecan, Switzerland). The hydrogels were washed 3 times with PBS, and 100 ul fresh media was added before the cells were placed back in the incubator. Viability was calculated by subtracting the signal of a respective hydrogel (HA-PEG and Matrigel) without encapsulated cells and normalized either to the percentage of the HA-PEG condition average or the percentage of day 1 average signal for the respective condition.

**Immunocytochemistry and imaging after spontaneous 3D differentiation:** After 7 days of spontaneous differentiation in hydrogels, cell media was removed, and the gels were washed with PBS before fixation with 10 % formalin solution (Sigma-Aldrich, USA) for 30 min at RT. After washing with PBS blocking buffer was added consisting of 10 % goat serum (Sigma-Aldrich, USA) and 0.1 % Triton-X in PBS. Doublecortin Antibody (Cell signaling, USA) was diluted 1:800 and left to incubate for 24 h at 4 °C. The primary antibody solution was removed, and the hydrogels were washed with PBS. Anti-rabbit CF633 (Sigma-Aldrich, USA) was diluted 1:1000, added to the wells and left to incubate overnight at 4°C, protected from light. The secondary antibody solution was removed, and the hydrogels were washed with PBS. Phalloidin CF®488 (Biotium, USA) was diluted at 1:40 and added to the wells to incubate for 90 min at RT. The solution was then removed, and the hydrogel was carefully washed with PBS, after which nuclear stain Hoechst was added at a dilution of 1:2000. After 10 min of incubation at RT, the solution was removed, and the hydrogels were washed with PBS. Antibodies, nuclear stain, and Phalloidin were diluted in a 1 % goat serum buffer and 0.1 %

Triton-X in PBS. The hydrogels were kept at 4 °C in PBS, protected from light until imaged. The hydrogels were imaged with a confocal scanning laser microscope Zeiss LSM 900 (ZEISS Research Microscopy Solutions, Germany)

**Image analysis:** Confocal images of differentiated lt-NES were composed using maximum projection, and brightness/contrast was adjusted to emphasize morphology. Two separate areas per hydrogel were chosen and imaged of each replicate, a total of twelve images of each hydrogel condition. One image of each condition was then presented in the paper through a blind test selection by an independent person.

**mRNA expression analysis:** The hydrogels were lysed and collected from the well plate and mechanically disrupted using a 27-G needle (Sigma Aldrich, USA), after which RNA was extracted using the RNeasy Mini Kit (Qiagen, Germany). RNA concentration was measured using MySpec (VWR, USA). cDNA synthesis was done with the High-capacity RNA-to-cDNA kit (Thermo Fisher, USA) on a thermal cycler (VWR, USA). The cDNA samples were incubated in Fast Advanced Master Mix together with gene-specific TaqMan PCR primers: GAPDH (Hs02758991_g1), SOX2 (Hs01053049_s1), NES (Hs04187831_g1), TUBB3 (Hs00964963_g1) and SYN1 (Hs00199577_m1), all from Thermo Fisher Scientific, USA. All samples were run on the BioRad CFX96 Touch Real-Time PCR Detection System using the multiplex option for superior $C_t$ quantification.

All samples were analyzed as technical duplicates multiplexing the gene of interest (GOI) and the housekeeping gene (HG) GAPDH. The expression of each GOI is presented as $\Delta\Delta Ct$ normalized to the expression of a reference sample (RS) of neuroepithelial stem cells using the following equations:

$$\Delta Ct = Ct(GOI) - Ct(HG)$$

$$\Delta\Delta Ct = \Delta Ct(RS) - \Delta Ct(GOI)$$

Linear mixed models (LMM) were used to calculate the significance of the different hydrogel conditions normalized to the RS in Origin Pro. Samples with a $Ct$ higher than 35 and duplicates with a variation higher than one were excluded from the analysis.

**3D bioprinting:** 3D bioprinting of SH-SY5Y laden hydrogels was conducted on a multi-nozzle Cellink BioX (Cellink AB, Sweden). The bioink was prepared in the same manner as described in the formation of hydrogels. The components were suspended in PBS before use. The bioink was loaded into a 3 ml syringe fitted with a sterile blunt needle (27G, 0.2 mm inner diameter). Single-layer square hydrogels (1 × 1 cm) was extruded at a rate of 17 mm $s^{-1}$ and 1 kPa pressure at room temperature with different extrusion patterns and density of the filaments of extrusion (infill density). Cell viability of undifferentiated SH-SY5Y cells was performed 24 hours after bioprinting using LIVE/DEAD Cell Viability Assay (Thermo Fisher Scientific, USA). Briefly, a solution of culture grade PBS with 2 µM calcein AM and 4 µM ethidium homodimer-1 (EthD-1) was added to the printed structures and incubated at 37°C for 30 min. Following the incubation, the samples were imaged using confocal microscopy (Zeiss LSM 780, ZEISS Research Microscopy Solutions, Germany) at excitation wavelengths of 488 nm (calcein AM – Live) and 543 nm (EthD-1 – Dead), respectively. Cell counts were performed manually from the obtained images. A 400 µm z-stack with 20 slices was imaged using 10 X objective (N=3 per group). The 3D image was converted to a 2D projection to quantify the cell viability using Image J Software.

**Syringe Ejection of NES in HA:PEG and media:** lt-NES were thawed and spun down in differentiation media. The cells were then resuspended at a 3.3 million cells/ml concentration in either 0.724:0.276 HA-BCN:$(PEG\text{-}Az)_8$ or DMEM/F12+Glutamax supplemented with 1 % N2 0.1 % B27 supplement and loaded into a 1 ml syringe with a 27-G needle attached. To optimize the procedure, a 5 mg/ml collagen gel, gelation at 37°C, PureCol EZ gel (Advanced BioMatrix, USA) were used. A volume of 30 µl cell suspension per well was ejected into either a half area 96 wp (hydrogel condition) or into a 48 wp (media condition) (WVR, USA) using an Aladdin Single-Syringe infusion pump at a rate of 50 µl/min. Cell viability was assessed immediately after ejection and after 24 h, during which the ejected cells were kept in an incubator until the assay. LIVE/DEAD® Viability/Cytotoxicity kit was used from which a solution of 2 µM calcein AM and 4 µM EthD-1 was prepared in PBS. The solution was added on top of the ejected cells and left to incubate for 40 min. The cells were then imaged with confocal microscopy (Zeiss LSM 700, ZEISS Research Microscopy Solutions, Germany) at excitation 488 nm (calcein AM – Live) and 543 nm EthD-1 – Dead) respectively. Each hydrogel was imaged with three stacked slices, and the media condition wells were imaged at three separate regions each. Cell counts were performed in Fiji.

**Statistical analysis:**

The statistical analysis for the viability of 3D-differentiation of lt-NES, mRNA expression and survival of ejected lt-NES was performed using Origin Pro (OriginLab, USA). P-values were derived using linear mixed models (LMM). In addition, statistical analysis of Alamar blue measurements and rheological storage modulus was performed, and p-values derived using one-way ANOVA with Tukey's HSD. No significance is indicated if the p-value is <0.05.

**Supplementary Figures**

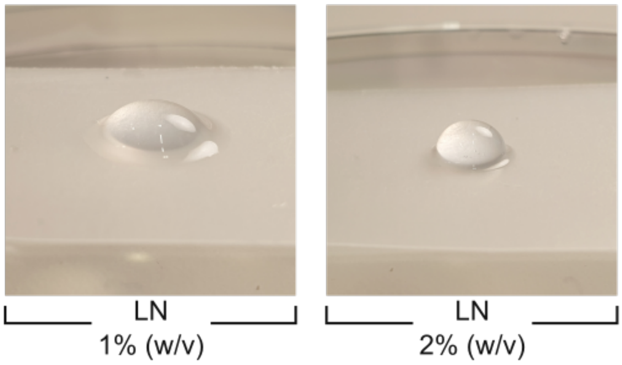


**Figure S1.** Photos of HA-PEG hydrogels functionalized with LN-(p)-Az.

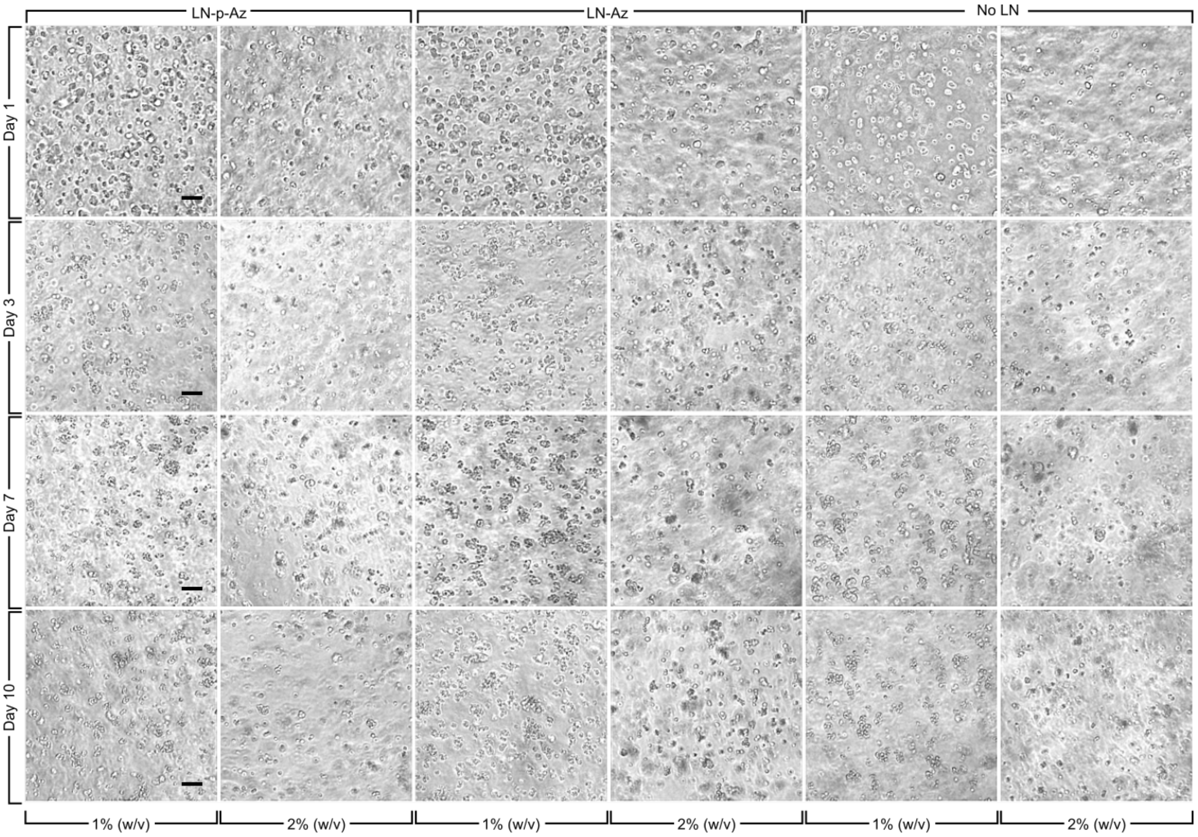


**Figure S2.** Undifferentiated SH-SY5Y cells cultured in the HA:PEG hydrogels with/without LN. Scale bars: 200 µm.

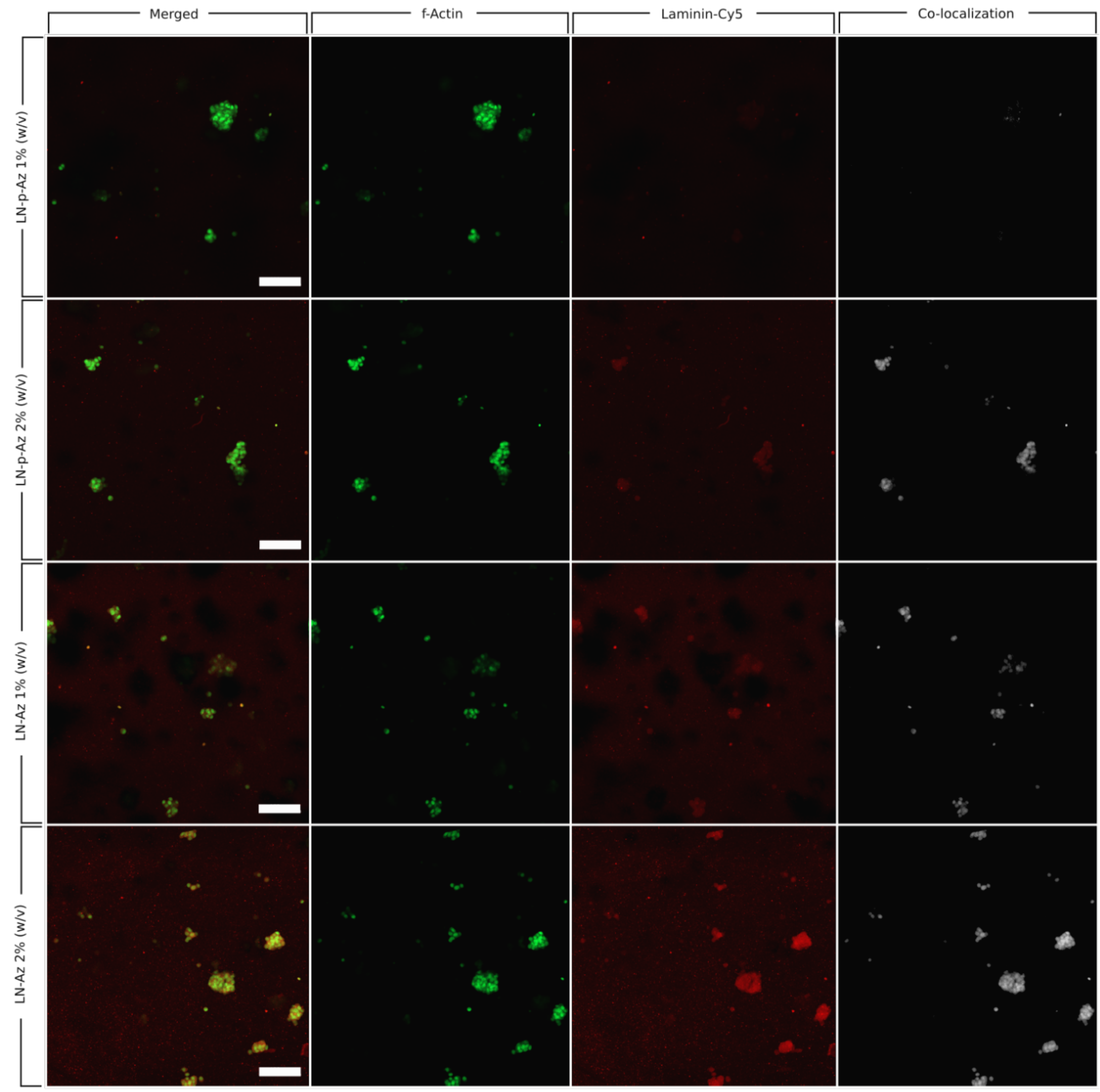


**Figure S3.** Confocal images of undifferentiated SH-SY5Y cells cultured in the HA:PEG hydrogels with/without LN. Green (actin), red (Cy5-labeled LN). Scale bars: 100 µm.

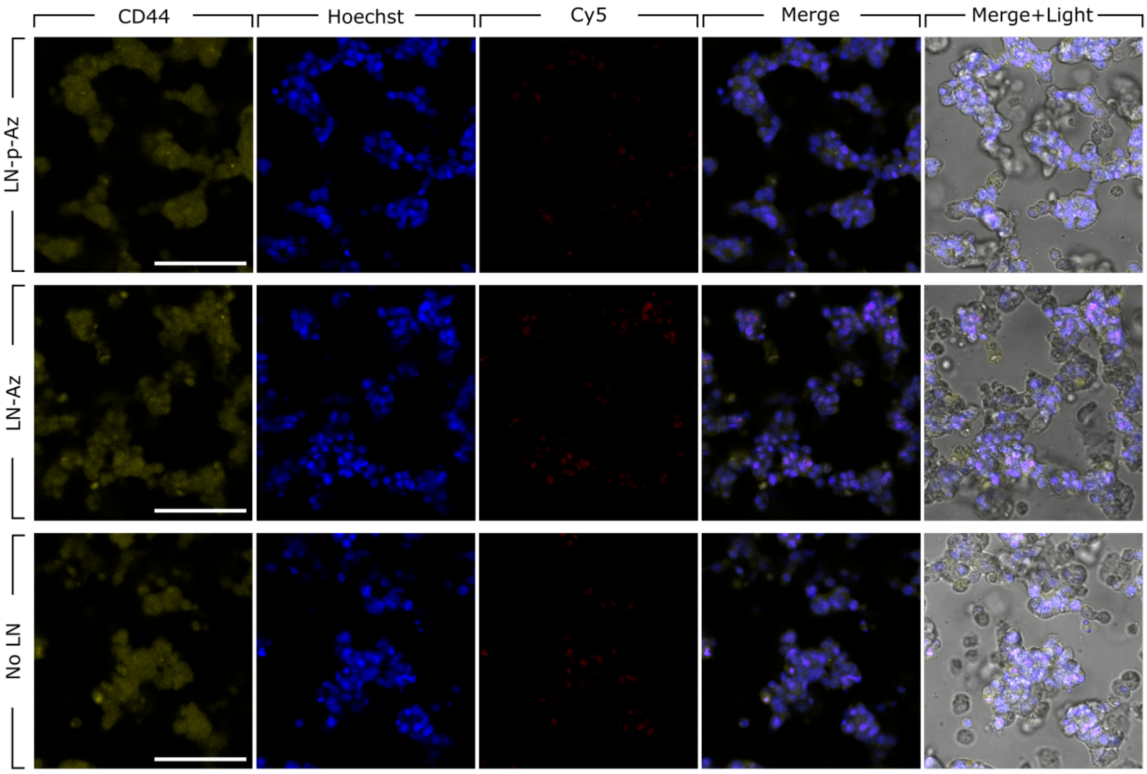


**Figure S4.** CD44 immunostaining of undifferentiated SH-SY5Y HA:PEG hydrogels with and without conjugated LN (Day 7). Scale bars: 100 µm.

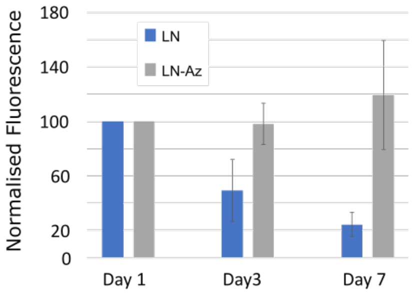


**Figure S5.** Viability of differentiated SH-SY5Y cells in HA:PEG-LN and in HA:PEG hydrogels supplemented with comparable concentrations of non-conjugated LN, N=4 for each condition.

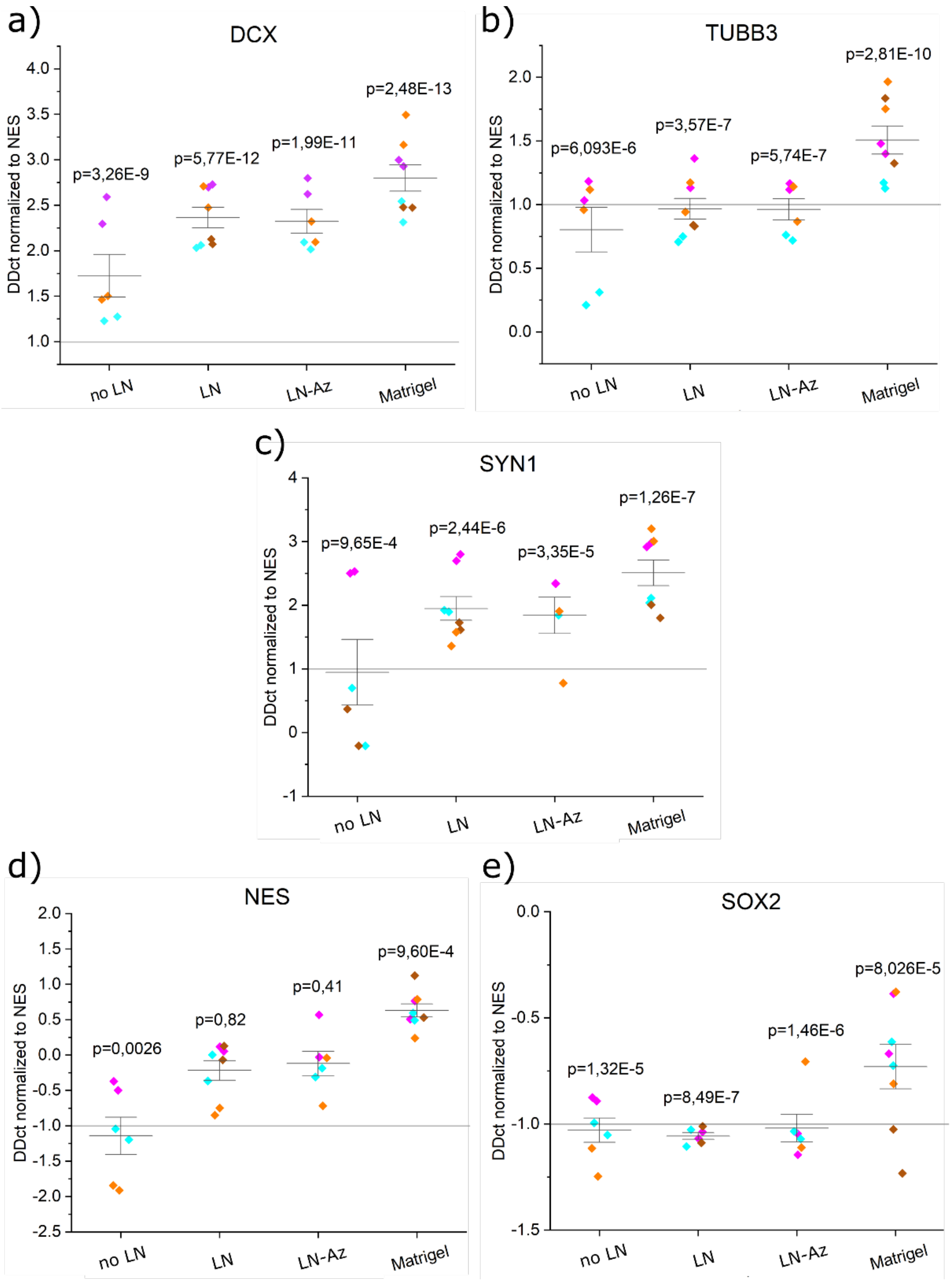


**Figure S6**. mRNA expression analysis of spontaneous 3D-differentiation in different hydrogel conditions. a) DCX, b) TUBB3, c) SYN1, d) NES, and e) SOX2. All samples contain the housekeeping gene GAPDH. P-values were derived with LMM in Origin Pro. Data is normalized to lt-NES. Data was collected as duplicates from three or four independent experiments.

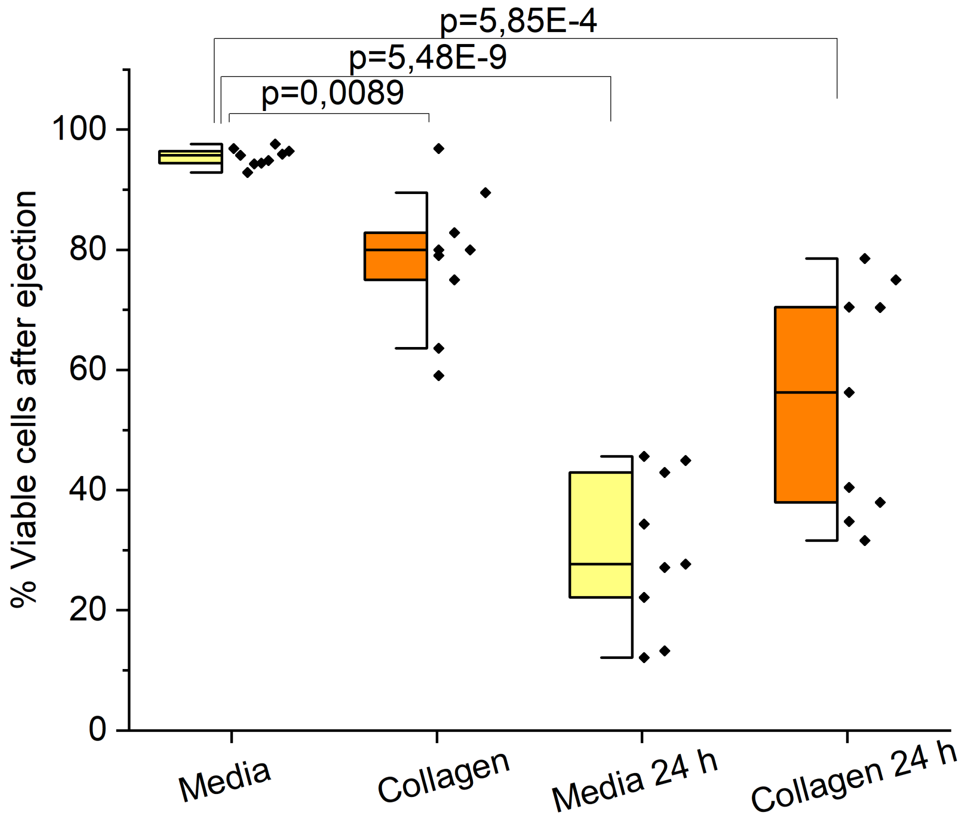


**Figure S7.** Survival of ejected lt-NES in collagen and cell media. Data was collected from one individual experiment, N=9, where N represents one ejected replicate. No data were excluded. P-values were derived using LMM in Origin Pro.

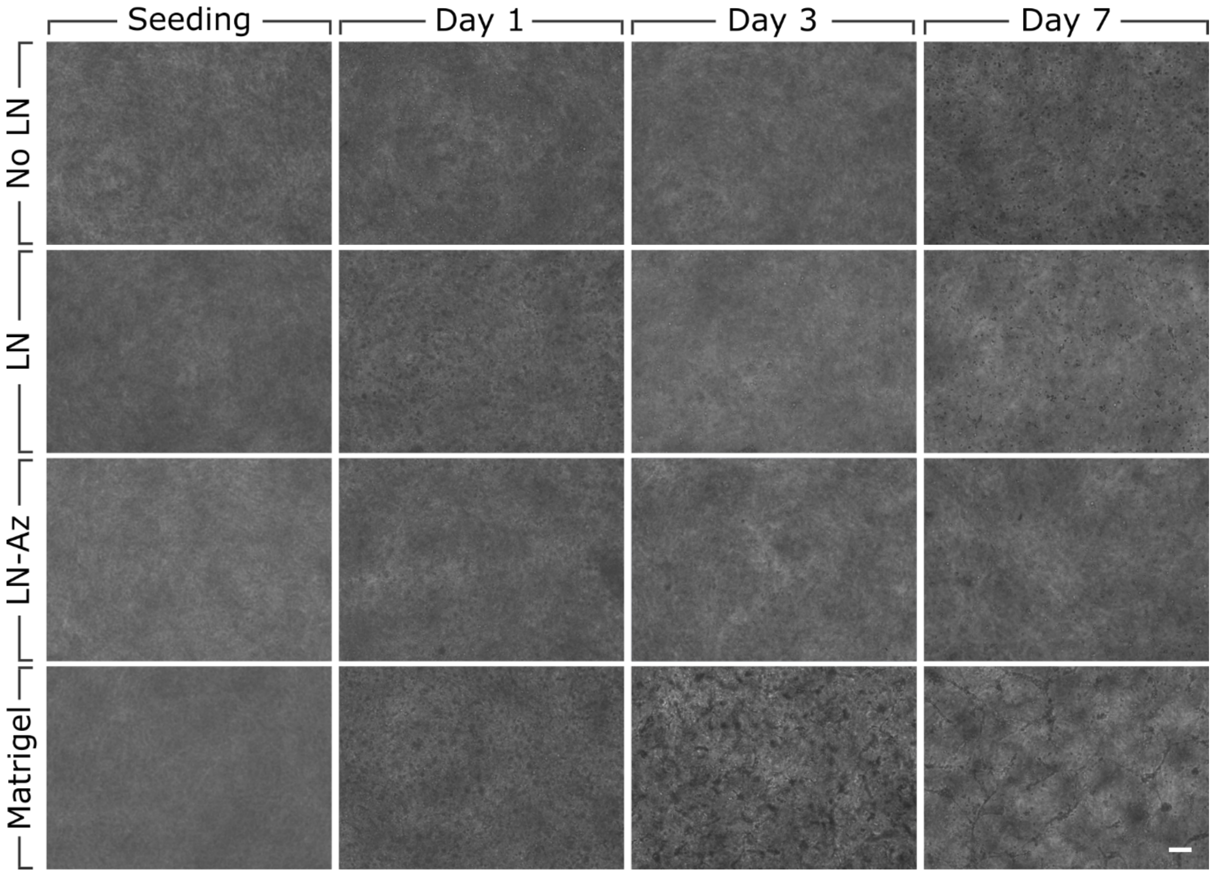


**Figure S8.** Brightfield images of spontaneous 3D-differentiation in respective hydrogels. Scale bar: 100 μm.

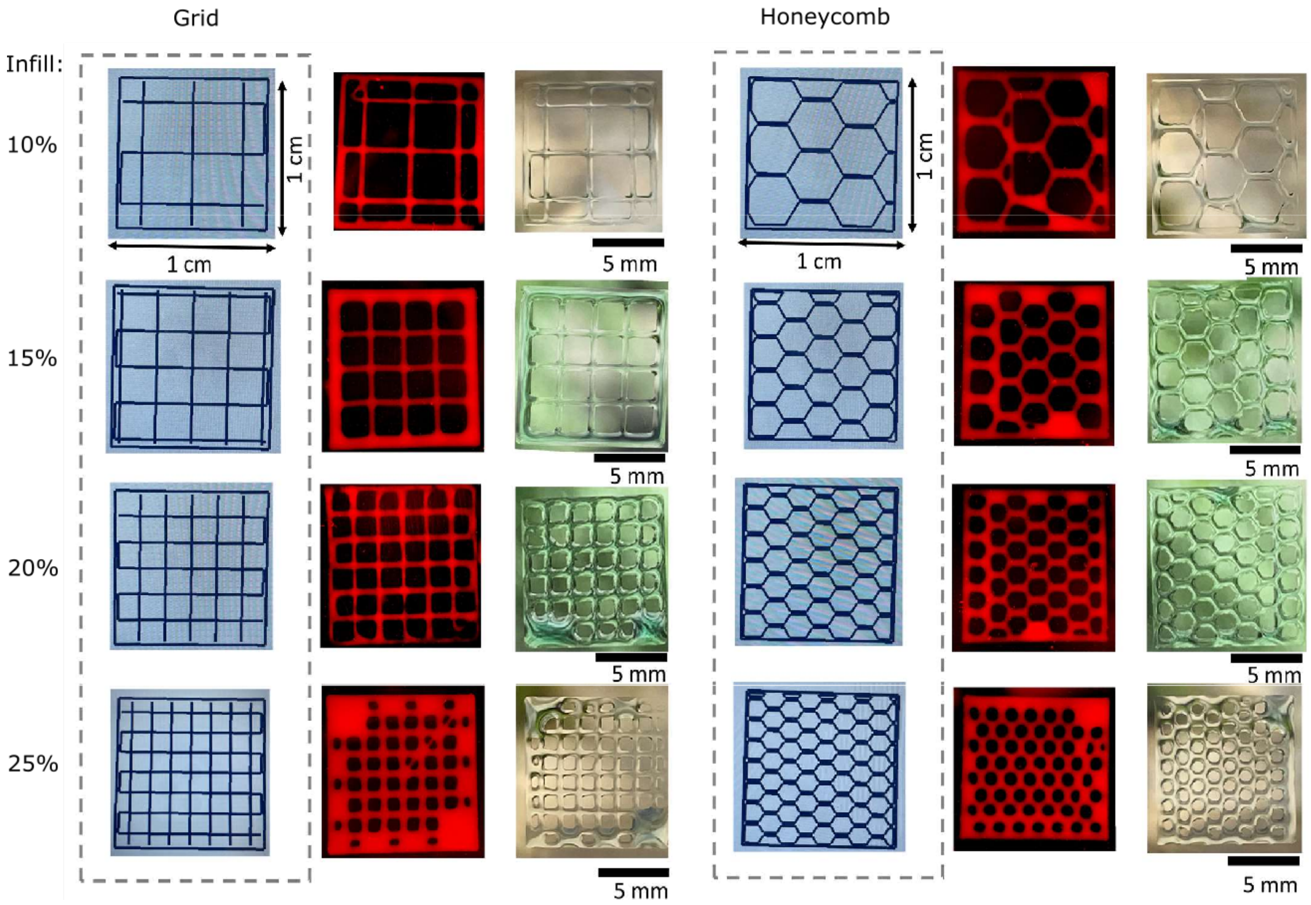


**Figure S9.** 3D bioprinting resolution as a function of infill for a grid (left) and a honeycomb structure.
